# Multi-kingdom quantitation reveals distinct ecological drivers of predictable early-life microbiome assembly

**DOI:** 10.1101/2020.03.02.970061

**Authors:** Chitong Rao, Katharine Z. Coyte, Wayne Bainter, Raif S. Geha, Camilia R. Martin, Seth Rakoff-Nahoum

## Abstract

The infant gut microbiota develops remarkably predictably^1–7^, with pioneer species colonizing after birth, followed by an ordered succession of other microbes. This predictable assembly is vital to health^8,9^, yet the forces underlying it remain unknown. The environment, host and microbe-microbe interactions are all likely to shape microbiota dynamics, but in such a complex ecosystem identifying the specific role of any individual factor has remained a major challenge^10–14^. Here we use multi-kingdom absolute abundance quantitation, ecological modelling, and experimental validation to overcome this challenge. We quantify the absolute bacterial, fungal, and archaeal dynamics in a longitudinal cohort of 178 preterm infants. We uncover with exquisite precision microbial blooms and extinctions, and reveal an inverse correlation between bacterial and fungal loads the infant gut. We infer computationally then demonstrate experimentally *in vitro* and *in vivo* that predictable assembly dynamics are driven by directed, context-dependent interactions between microbes. Mirroring the dynamics of macroscopic ecosystems^15–17^, a late-arriving member,*Klebsiella,* exploits the pioneer,*Staphylococcus,* to gain a foothold within the gut. Remarkably, we find that interactions between kingdoms drive assembly, with a single fungal species,*Candida albicans*, inhibiting multiple dominant gut bacteria. Our work unveils the centrality of simple microbe-microbe interactions in shaping host-associated microbiota, a critical step towards targeted microbiota engineering.

---

Humans are colonized by vast communities of microbes, particularly within the gastrointestinal tracts, that play key roles in host health^8,9^. However, we do not start life with these microbiota. Instead, infants are generally born uninhabited and their gut microbiota gradually assembles after birth^1–7^. Remarkably, this developmental process occurs in a strikingly predictable manner, with specific bacterial taxa establishing at distinct points in infant life^18–22^. Establishing the pattern of early life microbiome development is critical to infant and later-life health, with disruptions to microbiota development linked to a range of disease outcomes, morbidity and mortality, particularly within vulnerable preterm infants^1,23–27^. Yet despite their apparent importance, we do not understand what drives these patterned progressions^11–13^. Host physiology, diet, antibiotics, and the interactions between individual microbes are all likely to influence microbiota development. But with so many moving parts, the specific role of any individual factor has remained unclear. We do not know whether this predictable assembly simply reflects order of exposure, or changes in infant diet, or whether – as in macroscopic ecosystems – pairwise interactions between individual taxa integrate to shape the community as a whole^15–17^. Indeed, disentangling how and why microbial communities change over time remains a major challenge not just for the human microbiota, but for host-associated and environmental microbiomes more broadly.

Our ability to identify causal drivers of microbiota development has been held back not just by the intrinsic complexity of microbial ecosystems, but also by fundamental limitations in how we quantify community composition^10–13^. First, while next-generation sequencing (NGS) has provided a comprehensive map of bacterial diversity within the human gut^28,29^, we still know little of the other microorganisms, such as fungi and archaea, that colonize the infant microbiota^30–32^, constraining our ability to identify inter-kingdom microbial interactions that drive ecosystem dynamics^33^. Second, NGS data typically chart only the relative abundances of taxa – that is, they quantify the proportion of different microbes within a given community, but not their absolute amounts. If a species increases in relative abundance over time we cannot determine whether that species is blooming or simply others are dying out. The compositional nature of relative abundance data can thus mask true community dynamics (Fig 1a), critically undermining our ability to identify the biotic and abiotic forces that shape microbiota change^34–36^. Here we used a scalable multi-kingdom quantitation method to map absolute microbiota dynamics in a longitudinal cohort of preterm infants. Remarkably, by fitting these data to a comprehensive ecological model we demonstrated that within and between-kingdom microbial competition and exploitation shape the predictability of early-life microbiome assembly.

**Figure 1.**
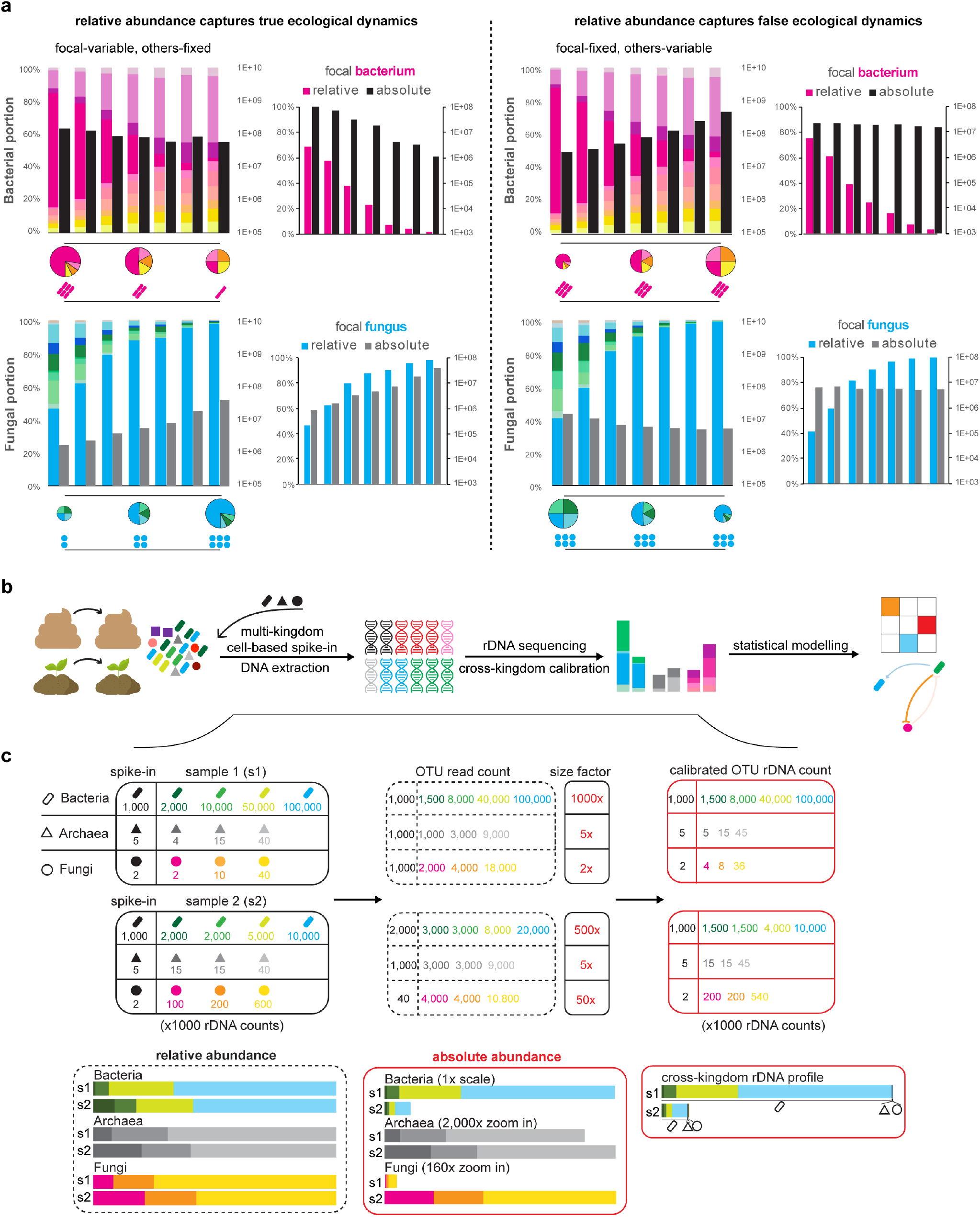
Multiple Kingdom SpikeSeq enables robust and reliable quantitation of absolute abundances.

**a**, Absolute abundance data disentangle two distinct mock ecological dynamics. Two sets of defined consortia were assembled to simulate a “true” (left) and a “false” (right) negative interaction between one focal bacterium and one focal fungus, by varying the abundances of either these focal species or other background members. The quantitation of focal species highlights the distinct patterns and conclusions of cross-kingdom interaction derived using relative versus absolute abundance data. **b**, Overview of the multi-kingdom quantitation pipeline to deconstruct mechanisms of microbiome dynamics. Prior to DNA extraction, defined amounts of multi-kingdom spike-in cells are added to human or environmental longitudinal samples as internal controls. Absolute abundance-based multi-kingdom dynamics are then characterized through standard rDNA amplicon sequencing. Further statistical modelling of these dynamics and relevant metadata predicts underlying microbe-microbe and microbe-host/environment interactions. **c**, Schematic example of MK-SpikeSeq. Two hypothetical samples with known abundances of each taxon are subjected to MK-SpikeSeq to generate cross-kingdom community profiles. The size factors used in back-normalization are derived by the known abundances of spike-in species. The relative abundance profiles without normalization give different and misleading conclusions compared with the calibrated absolute abundance profiles. MK-SpikeSeq provides extra information to compare across kingdoms in each sample.

## A scalable NGS pipeline to quantify multi-kingdom absolute abundances

To identify causal drivers of change within any microbial community one must quantify the absolute changes in the range of biotic community members over time. To achieve this, we developed a cell-based multiple kingdom spike-in method (MK-SpikeSeq) that quantifies the absolute abundances of bacteria, fungi and archaea simultaneously within any given microbiome (Fig 1b). Specifically, we add to each sample defined numbers of exogenous microbial cells of each kingdom and perform kingdom-specific rDNA amplicon sequencing to obtain relative abundances in each kingdom. The spiked-in cells serve as internal controls for the entire sample processing, including DNA extraction and library preparation. As abundances of the spike-in cells are known, we can back-normalize and calculate absolute abundances of all community members, across all three kingdoms (Fig 1c). As our primary objective here is to study the mammalian microbiota, our spike-in contains the bacterium *Salinibacter ruber*^37^, the fungus *Trichoderma reesei* and the archaeon *Haloarcula hispanica*, each selected based on their absence or rarity in mammalian microbiomes (Supplemental Table 2). Crucially though, our method can be easily adapted via spike-in choice and abundance to target any host-associated or environmental microbiome, and can be combined with shotgun metagenomics to capture viral members and enable strain-level absolute quantification.

We tested the ability of MK-SpikeSeq to sensitively and specifically measure absolute abundances in comparison to existing methods^35–38^ for quantifying microbial loads in diverse microbiomes. Total DNA quantification cannot distinguish between different microbial kingdoms (Supplemental Fig 1a). Similarly, cytometry-based imaging^36^ cannot readily distinguish archaea from bacteria, nor fungi from other eukaryotes (such as mammalian cells) due to similar scatter properties, as exemplified in samples that have high abundances of archaea, fungi, and intestinal epithelial cells (Supplemental Fig 1b). Quantitative PCR (qPCR) and DNA-based spike-in methods are able to target specific kingdoms^38^, but are vulnerable to underestimation of microbial abundances in samples with low microbial loads or high host cell contamination (Supplemental Fig 1c-e) – a particular problem when studying inflammatory diseases, where fecal samples can contain millions of inflammatory or epithelial cells per gram^39^. In contrast, MK-SpikeSeq can readily quantify each kingdom and is robust to variable DNA extraction efficiency (Supplemental Fig 1c-e). MK-SpikeSeq can, therefore, generate highly sensitive, robust and scalable absolute abundance measurements for individual taxa across multiple kingdoms within any given microbiome (Fig 1a, Supplemental Fig 2) – a key requisite for identifying true community dynamics in microbiota with unknown membership.

## Measuring multi-kingdom absolute abundances in the infant gut

Having developed and validated MK-SpikeSeq, we then built a high-resolution multi-kingdom picture of infant microbiota dynamics. Specifically, we assembled a prospective cohort of 178 preterm infants from a tertiary-care neonatal intensive care unit (NICU). We focused on preterm infants due to their clinical relevance and because they are amenable to high-frequency longitudinal sampling with readily available clinical and dietary metadata. These features render the preterm gut both an important and a tractable system for establishing a proof of principle understanding of microbiota assembly. We sampled each infant within our cohort on approximately their first, 14th, 28th, and 42nd day of life, and for 13 infants we gathered nearly-daily stools for their first 6 weeks of life (940 samples in total). Together, this cohort enabled us to build a high-resolution quantitative atlas of microbiota development within the preterm infant gut.

Consistent with previous preterm gut studies^18–23^, we observed that preterm infant gut bacterial communities cluster primarily into four distinct community states. These community clusters were independent of diet or delivery mode (Supplemental Fig 3), and were characterized by the domination of one of four genera: *Staphylococcus, Klebsiella, Escherichia* or *Enterococcus* (Fig 2a). Importantly, the bacterial community within our preterm cohort, as observed in numerous cohorts across continents^18–23^, developed in a predictable and highly dynamic manner over time. Most infants were initially dominated by *Staphylococcus,* then transitioned to a state dominated by one of the other three genera as infants aged (Fig 2a, b, Supplemental Fig 3), with total bacterial load gradually increasing over time (Supplemental Fig 6a). Indeed, we saw a similar positive correlation between archaeal abundance and infant age (Supplemental Fig 6b). Importantly, comparing the absolute and relative abundances of these dominant genera illustrated how compositional data can misattribute both how and when communities change. In one representative infant, relative abundances masked blooms of *Klebsiella* and *Escherichia*, and showed *Staphylococcus* and *Enterococcus* collapsing in the community when their abundances were, in fact, comparatively stable (Fig 2c).

**Figure 2.**
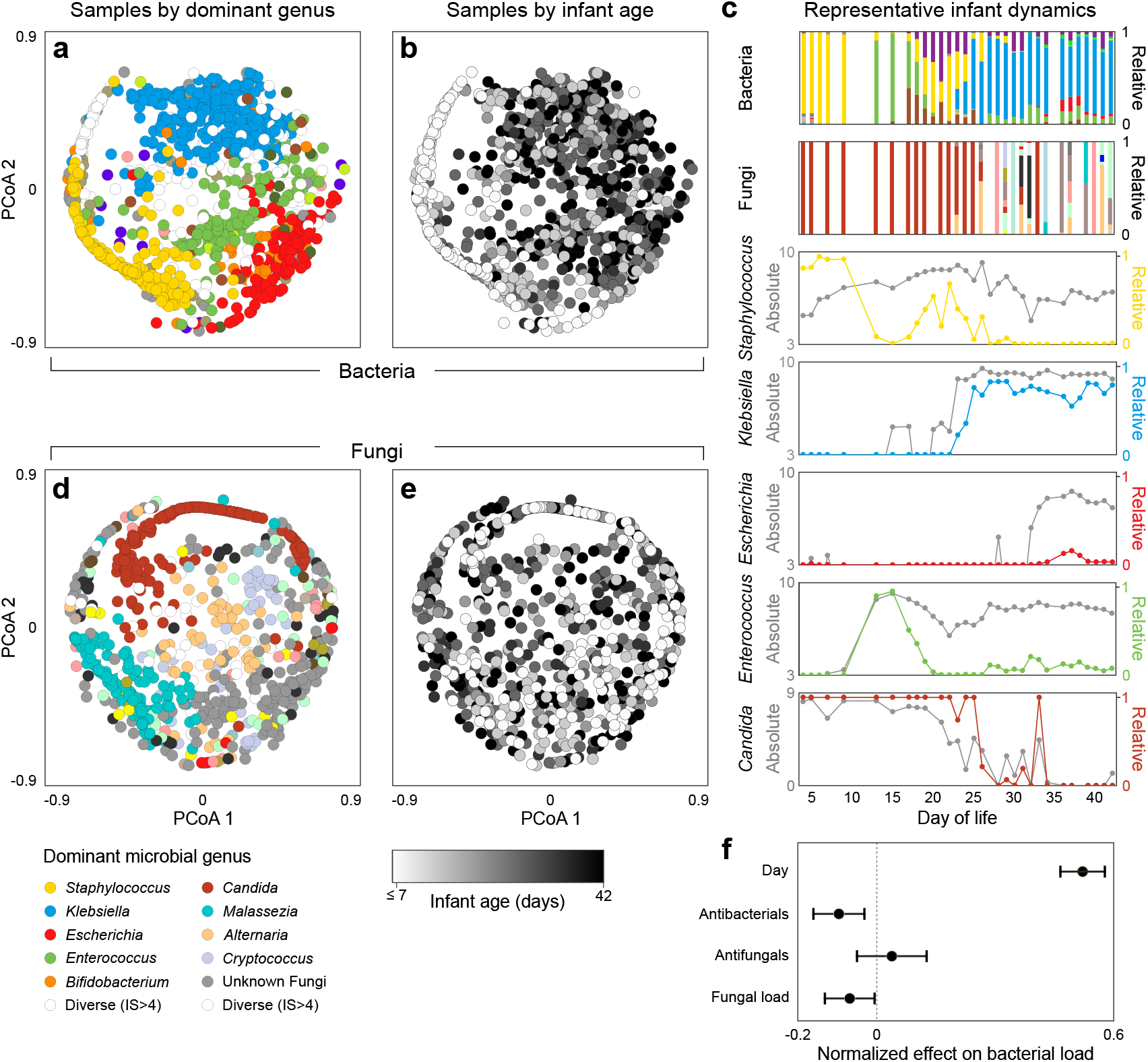
The preterm infant gut exhibits rich bacterial and fungal community dynamics. **a.** Principle Coordinate Analysis (PCoA) plot of Bray-Curtis dissimilarities between bacterial samples at the genus level. Each dot represents a sample, colored by the dominant genus present or white if diversity was high (Inverse Simpson index > 4). **b.**The same PCoA as panel **a** with samples instead colored by infant day of life, illustrating how bacterial community composition changes predictably over time. **c.** Microbiota dynamics of a single representative infant, highlighting the importance of gathering absolute abundances when studying microbiome ecology. Stacked bars represent total community composition (for full color schemes, see Supplemental Figs 3, 4). Line plots illustrate the relative (colored) and absolute (grey) abundances of individual genera. **d-e.** PCoA plots of fungal community composition colored by dominant genus (**d**) or infant age (**e**), indicating fungal community composition does not correlate with infant age. **f.** Effects of clinical and microbial factors on total bacterial load, quantified by a linear mixed effects model, suggesting a potential relationship between kingdoms within the preterm infant gut (error bars represent 95% confidence intervals). For panels **a**/**b**, number of samples n = 934, for **d**/**e** n=772, for **f** n=770.

In contrast to the predictable dynamics of bacterial communities, we uncovered diverse but unpredictable fungal communities within the preterm infant gut. On average, the fungal dynamics were noisier and exhibited far less temporal structure than bacterial communities (Supplemental Figure 5), with no clear correlation between fungal community composition or load and infant age (Fig 2d, e, Supplemental Figs 4, 6c). As with bacterial communities, we also observed instances where MK-SpikeSeq uncovered dramatic blooms and collapses in fungal genera masked by relative abundances (Figure 2c). However, though the fungal dynamics were themselves unpredictable, a linear mixed-effects model that accounted for infant age, antibacterials and antifungals, uncovered a significant negative correlation between bacterial and fungal loads (normalized effect size: −0.069, 95% CI: [−0.003, −0.134], Fig 2f). That is, when accounting for all clinical covariates, samples with higher fungal loads tended to have lower bacterial loads. This inverse relationship between bacterial and fungal communities suggested cross-kingdom interactions might be influencing preterm microbiota dynamics.

## Ecological inference reveals specific intra- and inter-kingdom interactions shaping infant microbiota assembly

Having gathered a high-resolution multi-kingdom picture of infant microbiota assembly, we next sought to identify causative factors driving the predictable community dynamics observed. To achieve this, we used Bayesian regularized regression to fit our longitudinal data to an extended generalized Lotka-Volterra (gLV) model – an approach only possible with absolute abundances. The gLV model assumes that the growth rate of an individual taxon depends upon the taxon’s intrinsic growth rate and interaction with kin, the effect of any antimicrobials present, and interactions between the focal taxon and other community members (Fig 3a; SI)^40–42^. Specifically, the model assumes microbes may interact with one another in a number of different ways, from bidirectional competition (−/−) such as when microbes compete for nutrients, to exploitation (+/−) wherein one microbe takes advantage of another, or not interact at all (0/0). The model also assumes each antimicrobial agent may inhibit, promote or have no effect on each community member. Together this yields a highly parameterised model of community dynamics. By fitting this model to our data using a highly conservative regularization framework we are powered to identify any microbe-microbe or microbe-antibiotic interactions playing a strong and consistent role in shaping community dynamics. This model enabled us to disentangle the effects of different biotic and abiotic interactions influencing microbiota assembly, independently of one another, missing community members (e.g. viruses), or any underlying host variability.

**Figure 3.**
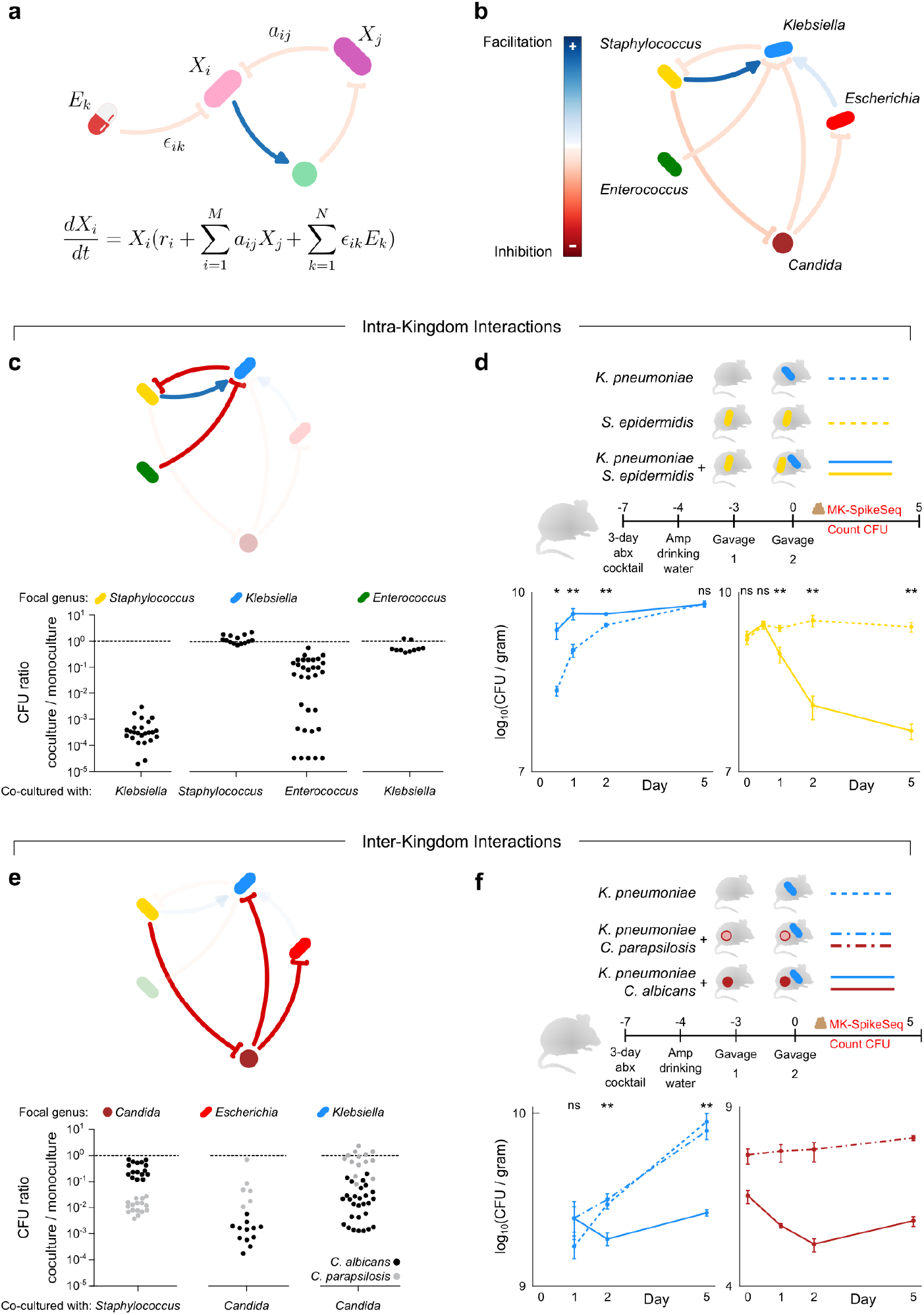
Pairwise intra- and inter-kingdom interactions drive predictable patterns of infant microbiome assembly. **a**, Schematic illustrating the generalized Lotka-Volterra (gLV) model used to identify causative drivers of microbiota dynamics. This model assumes the growth rate of each taxon, 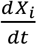 is determined by its own intrinsic growth rate *r_i_*, its interaction with other members of the community *a_ij_X_ij_*, and any environmental perturbation *ϵ_ik_E_k_*. **b,** The gLV model identified a network of within- and cross-kingdom interactions occurring between dominant members of the preterm infant gut that are predicted to affect microbiota dynamics. **c,** *in vitro* growth effects of infant isolates upon one another using monoculture and pairwise co-culture, testing predicted within-kingdom interactions. **d,** CFU counts quantifying microbial fitness *in vivo* in a specific pathogen free (SPF) mouse model, reproducing the predicted exploitation of *Staphylococcus* by *Klebsiella*. **e,** *in vitro* growth of infant isolates when growing alone and in pairwise co-cultures, testing predicted cross-kingdom interactions. Black dots indicate co-cultures with *C. albicans*, grey dots indicate co-cultures with *C. parapsilosis.* **f,** CFU counts quantifying microbial fitness *in vivo* in a SPF mouse model, reproducing the species-specific differences in the cross-kingdom inhibition observed *in vitro*. For panels **c/e**, each data point denotes one unique pair of strains of indicated genera. For panels **d**/**f**: *Klebsiella* was undetected at time 0 upon gavage; n=5 per group, error bars denote the standard error of the mean (SEM); * p<0.05, ** p<0.01, ns not significant, by student’s t test. For panel **f**, t test between *C. albicans* and *C. parapsilosis* groups.

This ecological inference suggested that strong intra- and inter-kingdom microbe-microbe interactions play a pivotal role in shaping microbiota assembly, independently of antibiotic perturbations (Fig 3b, Supplemental Figure 7). Remarkably, we inferred that the early colonizing *Staphylococcus* enhanced the growth of *Klebsiella* within the infant gut but was itself inhibited by *Klebsiella*. Thus our model suggests the characteristic transition from *Staphylococcus* to *Klebsiella* domination observed in preterm infants (Fig 2a, b) is shaped, in part, by *Klebsiella* exploiting the early colonizer to dominate the gut. We also found that *Klebsiella* itself was inhibited by another dominant genus *Enterococcus*, suggesting the distinct domination states of these two genera may be partly driven by one excluding the other. Perhaps most strikingly, consistent with the inverse correlation between bacterial and fungal loads, our analyses suggested that between-kingdom interactions play a key role in community dynamics. Specifically, we inferred that the fungal genus *Candida* inhibited both *Klebsiella* and *Escherichia*, but was itself inhibited by *Staphylococcus*. These results suggested that not only do preterm infants harbor diverse fungal communities, but that members of these communities are critically influencing larger scale community dynamics.

Notably, we discovered the overwhelming majority of interactions shaping preterm infant assembly are asymmetric, with commensalism (+/0), amensalism (−/0) and exploitation (+/−) together comprising over 80% of inferred microbe-microbe interactions (Supplemental Figure 7c). The importance of these directed, asymmetric interactions in shaping microbiota assembly not only sheds light on microbiota ecology – it also underlines the power of our absolute abundance-based inference to study microbiota communities. Without absolute abundances, ecological inferences are limited to correlational analyses. These analyses can identify positive or negative correlations between the frequency of taxa, but cannot determine directionally who is interacting with whom, nor identify asymmetric interactions^34,43–45^. Indeed, when applied to our dataset, correlational relative abundance analyses (FastSpar^46^) erroneously inferred that *Staphylococcus* inhibited *Klebsiella* and promoted *Candida* (Supplemental Fig 8). In other words, relative abundances and correlation analysis not only misrepresented the dynamics of infant microbiome assembly (Fig. 2), but also incorrectly misclassified the core ecological processes underlying these dynamics.

## *In vitro* and *in vivo* validation confirm microbial interactions shaping development

Our ecological inference indicated that microbe-microbe interactions are central to the predictable infant microbiome assembly. However, though we employed a highly conservative regularization framework to ensure robustness to spurious correlations, our predictions may still be vulnerable to unobserved confounding factors not explicitly incorporated in the model, such as diet, viruses, or the host itself. Indeed, though a small but increasing number of studies have used similar modelling approaches to infer interactions, very few predictions have been experimentally validated^40–42^. We therefore sought to determine whether we could reproduce our inferred interactions in a reductionist experimental system, focusing first on our predicted within-kingdom interactions. Specifically, we isolated *Staphylococcus*, *Klebsiella, Escherichia* and *Enterococcus* strains from five infants in our cohort, capturing several distinct species of each genus. We then performed monoculture and pairwise co-culture of these strains and used colony forming unit (CFU) yields to determine the pairwise fitness effects of strains upon one another. Remarkably, we were able to reproduce *in vitro* all of the inhibitory interactions predicted by our model, with growth effects largely conserved at the genus level (Fig 3c). *Klebsiella* strongly inhibited *Staphylococcus*, reducing *Staphylococcus* yields by over 1000-fold, while *Enterococcus* variably but consistently inhibited *Klebsiella* growth (Fig 3c). Importantly, as predicted, *Klebsiella* showed no effect on *Enterococcus* growth, confirming this interaction is amensalism as opposed to bi-directional competition – validating the specificity and directionality of our inference.

In contrast to our predictions that *Staphylococcus* benefitted *Klebsiella* during infant microbiome assembly, we did not observe a positive effect of *Staphylococcus* on *Klebsiella* growth *in vitro* (Fig 3c). Given the strength of the predicted *Klebsiella-Staphylococcus* exploitation, we hypothesized that this interaction may be context dependent, i.e., *Klebsiella* only gains a benefit from *Staphylococcus* within the gut. To investigate this, we used two co-resident NICU isolates, *Klebsiella pneumonia* and *Staphylococcus epidermidis*, to test if *Klebsiella* benefited from the presence of *Staphylococcus in vivo* in the mammalian gut using a mouse model of intestinal colonization (Fig 3d). Specifically, we used MK-SpikeSeq and CFU counts to measure the fitness of *K. pneumoniae* after gavage in mice pre-colonized with or without *S. epidermidis* (Fig 3d, Supplemental Fig 9a). Strikingly, we observed that, as predicted from our modelling, *S. epidermidis* significantly enhanced the ability of *K. pneumoniae* to colonize the mouse gut, with *K. pneumoniae* exhibiting faster colonization if the gut was pre-colonized with *S. epidermidis* (Fig. 3d). Moreover, mice colonized with *K. pneumoniae* had significantly reduced levels of *S. epidermidis* than those without *K. pneumoniae*, with *S. epidermidis* declining alongside the rise in *K. pneumoniae* (Fig 3d). These *in vivo* data recapitulated the dynamics observed in infant assembly (Fig 2c) and demonstrated that the predictable patterns in infant microbiome assembly are due to exploitation of an early pioneer species by a late colonizer, establishing a striking parallel between human early-life microbiota development and macroscopic ecological succession.

Having validated our inferred within-kingdom interactions, we sought to validate the network of between-kingdom interactions predicted to influence preterm gut assembly (Fig 3e). Remarkably, each of our predicted between-kingdom interactions could also be reproduced within our *in vitro* system (Fig 3e). As predicted, *Candida* members caused on average a ~100-1000-fold inhibition of each *Enterobacteriaceae* and experienced a ~10-100-fold reduction in growth when co-cultured with *Staphylococcus*. Notably though, we observed a clear bimodal distribution in the strength of inhibition of different *Candida* isolates by *Staphylococcus* (Fig 3e), with the two levels corresponding to two *Candida* species, *C. albicans* and *C. parapsilosis.* Moreover, these two species also exerted differing inhibitory effects on *Enterobacteriaceae*, such that overall *C. albicans* both resisted *Staphylococcus* and inhibited *Enterobacteriaceae* more than *C. parapsilosis* (Fig 3e). To examine if this species-specific inhibition of *Enterobacteriaceae* by *Candida* occurred *in vivo*, we pre-colonized mice with either *C. albicans*, *C. parapsilosis* or vehicle control, then introduced *K. pneumoniae* and measured microbial colonization dynamics. We observed a significantly reduced colonization of *K. pneumoniae* in the presence of *C. albicans*, compared to vehicle-control or pre-colonization with *C. parapsilosis*, validating both the species-specificity in the fungi-bacteria interaction and its occurrence within the mammalian gut (Fig 3f, Supplemental Fig 9b). Together, our data demonstrated a novel species-specific cross-kingdom interaction that shapes the preterm infant gut microbiota.

The *de novo* assembly of the infant gut microbiome is remarkably ordered, with pioneer microbes colonizing first, followed by predictable waves of other microbes. To date, the forces driving these predictable transitions occur have remained elusive. Factors such as priority effects, diet and antibiotics, and the developing immune system are all thought to impact microbiota dynamics. However, with multiple interacting factors at play, disentangling the role of any individual process has remained a formidable task. Here we demonstrate the power of combining multi-kingdom absolute abundance quantitation, ecological modelling, and experimental validation to overcome this challenge. We have demonstrated that the predictable patterns of pre-term infant gut assembly are driven by direct, context-independent interactions between microbes. Just as in macroscopic ecosystems^15–17^, microbes exploit one another to establish within the infant gut, and direct interactions between kingdoms play a central role. The reducibility of gut microbiota assembly to simple, pair-wise interactions has profound implications for understanding and ultimately manipulating gut microbial ecosystems in health and disease.

## Materials and Methods (Supplemental information)

### Section 1 – MK-SpikeSeq development and validation

#### MK-SpikeSeq development

The backbone of MK-SpikeSeq is the addition of a defined number of bacterial, fungal and archaeal cells to the sample of interest. These cells serve as internal controls for sample processing and subsequent normalization. In this work we developed the MK-SpikeSeq method primarily for use on mammalian stool samples, however, our method can easily be adjusted for other samples types by changing the spike-in taxa and the quantity added. The criteria for spike-in usage are:

1. The spike-in taxa should absent in the sample
2. Once added, the spike-in should account for 0.1%~10% of the sample quantity (assuming the sequencing depth is >10K per sample; higher depth of sequencing could yield broader range of detection).

To test these qualifications, multiple representative test samples (ideally covering both upper and lower bounds of absolute abundance) should be used for preliminary DNA extraction. The targeted ribosomal DNA (rDNA, e.g., 16S) of these test samples should be quantified using rDNA-specific quantitative PCR (qPCR, Supplemental Table 1) and literature reviewed (or empirically sequenced) to confirm the absence of the selected spike-in taxa. To identify an appropriate quantity of spike-in cells, 10-fold serial dilutions of a defined number of spike-in cells should be DNA extracted and qPCR quantified in the same way as the test samples to identify the dilution of spike-in that meets the 0.1%~10% abundance requirement for most test samples (see the stool example below).

For human and mouse stool samples, we selected *Salinibacter ruber* DSM 13855, *Haloarcula hispanica* ATCC 33960 and *Trichoderma reesei* ATCC 13631 to use as spike-in taxa for bacteria, archaea and fungi, respectively. These taxa we chosen based on their absence/rarity in a systematic screen of several major public depositories of host-associated microbiome data (Supplemental Table 2). They also have the advantage of being publicly available and having completed genomes. Spike-in cells were grown according to supplier’s instructions: *S. ruber* and *H. hispanica* were cultured in ATCC medium 1270 and *T. reesei* was grown on Potato Dextrose Agar medium. These bacterial cells and fungal spores were harvested and homogenized in DNA-free PBS, passed through 70 um cell strainers and quantified based on optical density unit (ODU) at 600nm. We empirically determined to spike in 10^−2 ODU *S. ruber*, 10^−5 ODU *H. hispanica* and 10^−4 ODU *T. reesei* to each 100 mg human stool or 10 mg mouse stool based on:

1. 10^−2 ODU *S. ruber* accounted for on average ~1% of the bacterial 16S abundances of stool samples, and
2. 10^−5 ODU *H. hispanica* and 10^−4 ODU *T. reesei* were low enough to complement the usually low archaeal and fungal signal in stools, but high enough for robust pipetting of consistent cells numbers (>10^3 cells).

To implement our spike-in protocol, stool samples were first weighed and aliquoted into bead-beating DNA extraction tubes. Total amounts of spike-in cells were adjusted as per weight of samples, and the spike-in volumes to be added into the samples prior to DNA extraction were calculated based on the measured densities of spike-in cells. We used ZymoBIOMICS DNA extraction kit (Zymo Research, Irvine, California) for DNA extraction and performed standard two-step PCR amplification to construct rDNA amplicon libraries for Illumina sequencing. We empirically determined the rDNA abundance of each spike-in at the pre-determined level, using rDNA-specific qPCR against plasmid DNA standards that were constructed to contain tandem single copy of each rDNA. We measured the 10^−2/10^−5/10^−4 ODU multi-kingdom spike-in as: bac16S_Srub : arch16S_Hhis : ITS1_Tree ~ 10,000,000 : 50,000 : 20,000. Note that these cross-kingdom ratios were experimentally quantified but the exact numbers were arbitrary; as practically DNA extraction is not 100% efficient, the exact rDNA copy number per sample is less meaningful than the relative changes between samples in one study. These spike-in rDNA abundances were used as scaling factors in the later cross-kingdom normalization: Normalized OTU abundance = OTU read count / Spike-in read count * kingdom-specific scaling factor. See https://github.com/katcoyte/MK-SpikeSeq for our detailed MK-SpikeSeq protocol.

#### MK-SpikeSeq validation

We validated our MK-SpikeSeq method in two ways: 1) comparison with other existing approaches for absolute quantification, and 2) benchmarking using a series of defined mock samples.

To compare MK-SpikeSeq with other methods, namely total DNA yield, flow cytometric cell counting and rDNA-specific qPCR, we assembled a set of 40 test samples including human stools and soil samples (Supplemental Table 3). Each sample was homogenised (each 100 mg sample in 1 mL sterile PBS) by vortexing for 5 min, passed through 70 um cell strainers and then divided equally into three aliquots for independent use in flow cytometric cell counting, DNA extraction without spike-in and MK-SpikeSeq. For cell counting, we diluted samples further by 10 fold in 2% paraformaldehyde. We used the Bacteria Counting Kit (Invitrogen, Cat B7277) for cell staining and enumeration on a LSRFortessa 3-laser cell analyzer (BD Biosciences, San Jose, CA), and applied different flow settings and FSC/FITC gating strategies for prokaryotes and fungi, pre-determined using axenic bacterial and fungal cultures as positive controls. While prokaryotic gating was relatively robust against background noise, we found it was difficult to identify a gating specific for fungi against background (e.g., food debris and host cells). As a result, fungal cell counts were consistently over-estimated using three different gating strategies (Supplemental Table 3). For total DNA quantification, we performed technical duplicates for each sample using Quant-iT PicoGreen dsDNA Assay Kit (Invitrogen, Cat P7589) and excluded samples with concentration near or below detection limit (0.1 ng/ul) for correlational analysis (Supplemental Table 3). For qPCR, we performed technical triplicates for each sample and excluded samples with Ct > 30 or high variation (Ct SD > 0.05 * Ct Mean) for correlational analysis (Supplemental Table 3d). For MK-SpikeSeq, we used 10^−3 ODU *S. ruber*, 10^−5 ODU *H. hispanica* and 10^−4 ODU *T. reesei* for each 100 mg sample (lower *S. ruber* to accommodate to the lower bacterial density in soil samples). Systematic comparisons between these methods suggested that MK-SpikeSeq outperformed others in cross-kingdom sensitivity and specificity (see main text and Supplemental Fig 1a-c). We also performed rDNA amplicon sequencing of samples without spike-in, and demonstrated that exogenous spike-in did not alter rDNA relative abundance profiles of the communities (Supplemental Figure 1f).

To determine the sensitivity, robustness and accuracy of MK-SpikeSeq, we further generated multiple sets of defined mock samples. We first tested the limit of bacterial and fungal detection using serial dilutions of *Escherichia coli* (10^9 to 10^1 colony forming units CFU) and *Candida albicans* (10^7 to 10^1 CFU) axenic cultures, respectively. Because the range of detection using SpikeSeq is limited to the depth of sequencing, we examined two levels of spike-in: high spike-in (10^−3 ODU *S. ruber*, 10^−2 ODU *T. reesei*) targets high abundance samples, and low spike-in (10^−5 ODU *S. ruber*, 10^−4 ODU *T. reesei*) targets low abundance samples (Supplemental Table 4). We found that both spike-in levels could reliably detect a range of at least 6 orders of magnitude, and the low spike-in could successfully detected as low as 10 bacterial CFU within a sample (Supplemental Table 4, Supplemental Fig 1d). As a comparison, qPCR, the second-best method we benchmarked, failed to distinguish samples below 10^4 bacterial CFU (Supplemental Table 4, Supplemental Fig 1d).

We next determined the robustness of our method against background cells (e.g. host cells) using a set of samples with fixed amounts of *E. coli* and *C. albicans* (10^6 and 10^4 CFU, respectively) but variable amounts of host Caco-2 cells (up to 10^6 cells) in order to model conditions where > 10 million colonic cells are contained per gram stool. Using SpikeSeq we detected relatively consistent (<=2 fold difference) abundances of bacteria and fungi; however, qPCR undermeasured both bacteria and fungi by > 3 deltaCt (~10 fold) under the high host cell background (Supplemental Table 5, Supplemental Fig 1e).

Finally, we systematically examined the accuracy of MK-SpikeSeq in capturing community changes, using multiple series of defined mock communities (Supplemental Table 6). In the first two sample sets, we assembled single-kingdom communities composed of either ten bacteria or ten fungi, with fixed proportions of each community member. We then generated 10 instances of each community type with varying total abundances by serial dilutions. Our MK-SpikeSeq method reliably recapitulated the expected total abundances and compositions of each of these samples (Supplemental Fig 2a,b). In the third sample set, we mixed our defined bacterial and fungal communities in different ratios. Using MK-SpikeSeq, we correctly captured the bacteria-fungi ratios in these cross-kingdom samples (Supplemental Fig 2c). In the last two sample sets, we simulated a “true” and a “false” negative interaction between one focal bacterium (*Lactobacillus rhamnosus*) and one focal fungus (*C. albicans*), by changing the abundances of either these focal members or the abundances of other background members. Using relative abundance data, these two focal members showed undistinguishable inverse relationships in both sets; in contrast, absolute abundance data generated by MK-SpikeSeq faithfully disentangled these two fundamentally different relationships (Fig. 1c). Collectively, these data demonstrate that MK-SpikeSeq is sensitive for low-microbial abundance samples, robust to host cell background and accurate in revealing true cross-kingdom absolute abundances.

### Section 2 – Infant cohort: selection and analysis

#### Premature infant fecal sample biorepository

Study subjects and corresponding fecal samples were selected from a cohort of premature infants less than 33 weeks of gestation enrolled in the Infant Health Research Program at Beth Israel Deaconess Medical Center (BIDMC), Boston, MA from 2009 - 2013. The BIDMC Institutional Review Board (IRB) approved the collection of discarded specimens for the clinical biorepository after verbal consent from the infant’s parent(s) (IRB protocol number 2009P-000014). IRB approval was obtained for the analyses of these samples for this study under protocol 2017P-000632. All data were de-identified and available on request.

Infant were enrolled soon after admission to the neonatal intensive care unit (NICU) and every fecal sample was collected until six weeks postnatal age. All soiled diapers were placed in a 4 degree refrigerator. Fecal samples from all diapers representing a 24 hour period were sterilely scraped from the diapers, placed in a sterile cup, and homogenized by hand. Samples were then aliquoted in 1.9 mL cryovials and placed in a −80 degree freezer until sample analysis.

Maternal and infant clinical data were abstracted from the electronic medical record. The final cohort used for analysis was roughly equally split between male and female (92:86, M:F) and the average gestational age of each infant at birth was 29.6+2.5 weeks. The majority of infants within the cohort were born via Caesarean-section (131/178).

#### Sample collection and processing

We applied MK-SpikeSeq to the NICU cohort in two phases. To build a broad picture of preterm microbiota development during the first six weeks of life, we first collected a coarse timeseries of samples from 178 infants, sampling on approximately days 0, 14, 28, 42 of life. Overall, in this phase we collected 3.6±0.8 fecal samples per infant, yielding 631 samples in total. To build a higher resolution picture of microbiota development, we next selected 13 C-section infants (6 male, 7 female) for whom we collected every possible stool sample for the first six weeks of life, generating a total of 309 samples (on average 23.7±7.4 samples per infant). All samples were collected as raw stools and stored at −80C.

Following our established MK-SpikeSeq protocol (see earlier), we aliquoted ~50 mg of each stool sample into a bead-beating tube, added spike-in cells adjusted to the sample weight, then performed standard DNA extraction and amplicon library preparation in the 96-well format. We then sequenced samples, employing a different sequencing strategy in each phase. In the first phase, we performed Illumina Nextseq sequencing, featured by its shorter read (300-cycle, with read 1 extended to 200nt) but higher throughput (> 300M reads), to simultaneously profile bacterial (universal 16SV4), archaeal (archaeal-specific 16S) and fungal (ITS1) communities at (mostly) the family level in the 631 stool samples (Supplemental Tables 7-10). In the second phase, we took advantage of the long-read (600-cycle, 300PE) Miseq sequencing to obtain (mostly) genus level resolution of bacterial (16SV3V4) and fungal (ITS1) communities (after establishing the rarity of archaea in preterm infants) in the 309 stool samples (Supplemental Tables 7, 11-12).

To derive the OTU table for microbiota analyses, the obtained Illumina raw reads were demultiplexed, paired-end joined (only for Miseq 300PE reads), adapter trimmed, quality filtered (trimmed uniform to 100 nt for Nextseq reads only), dereplicated, denoised, and sequences mapped against public rDNA databases SILVA and UNITE. See https://github.com/katcoyte/MK-SpikeSeq for scripts used for Nextseq and Miseq data. Amplicon samples with < 1K total read counts or < 10 spike-in counts were excluded from further analyses (dropping 3 bacterial, 57 archaeal and 157 fungal samples in the Nextseq data, and 28 fungal samples in the Miseq data). For the archaeal OTU table, spurious bacterial OTUs based on taxonomic annotations were removed. Counts of spike-in OTUs (g__Salinibacter, g__Haloarcula, g__Trichoderma) were used in normalization for other taxa as described above.

All Illumina sequencing raw read, including cohort samples and validation samples, have been deposited at the European Nucleotide Archive (ENA) under study accession no. PRJEB36435.

#### Microbiota analyses and visualization

All data analyses and visualization were carried out in Python 3.7 or R. To visualize microbiota dynamics and group samples based on similarity we calculated the Bray-Curtis distance between each sample at the level of genus (*sklearn.metrics.pairwise.pairwise_distances(X, metric=’braycurtis’)*) then used multi-dimensional scaling to reduce our data to two dimensions (*manifold.MDS(n_components=2, eps=1e-12, dissimilarity="precomputed", max_iter=5000)*). We then identified clusters of similar samples using DBSCAN (*sklearn.cluster.DBSCAN(eps = 0.05, min_samples = 3)*). To determine whether microbiota clusters were associated with infant age, we performed a kruskal-wallis test on infant ages within clusters using the *kruskal* function from the *agricolae* package in R, with a bonferroni adjustment for multiple hypotheses. For all taxonomy-based visualization, following Taur et al, samples were classified as being “diverse” if they had an inverse simpson index > 4, and colored white. If not, samples were colored according to the dominant genus present.

For statistical analyses of total bacterial load, we combined our two sample groups to build a dataset of 169 unique infants for whom we had 770 total observations where both fungal and bacterial loads were accurately measured (170 samples from our original 940 were excluded due to either fungal or bacterial amplicon sequencing not achieving the requirement of a least 1K total read counts and 10 spike-in counts). We matched each sample to the number of antibiotic agents administered on the day of sampling, excluding any topical antibiotics (Bacitracin, Gentamycin Ophthalmic etc). We then fit a linear mixed effects model with baby ID as a random intercept, defined by the Wilkinson notation:

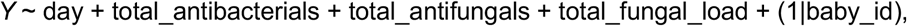

Where *Y* was the dependent variable, log_10_(total bacterial load) and the fixed effects covariates were day of life, the number of unique antibacterials or antifungals administered on each day, and log_10_(total fungal load). Notably, in any cases where total bacterial or fungal loads were measured as zero, we replaced these values with 10^3^ or 1 respectively. These values were our limits of detection for bacteria and fungi, with this change implemented to avoid underestimating loads when they fell below our detection window.

To calculate the noise profiles of bacterial and fungal dynamics, following Faust *et al*, we used the R package *seqtime* to first fit a spline to each ASV’s dynamics then quantify the noise profile for each prevalent ASV within each baby, using the function *identifyNoiseTypes*. We then compared the average proportion of black, brown, pink and white noise across babies for each kingdom, with black / brown noise indicating strong temporal structure and white noise indicated no temporal structure.

### Section 3 – Identifying drivers of microbiota development

To identify microbe-microbe and microbe-antimicrobial interactions shaping community dynamics we fit our data to an extended generalized Lotka-Volterra model (gLV) that accounts for antibiotic and antifungal usage. Under this framework, the growth rate of each microbial taxon *i* is expressed as,

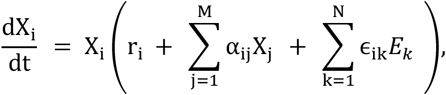

Where *X*_*i*_ is the abundance of taxon *i* and *r*_*i*_ is its intrinsic growth rate. *α_ij_* denotes the effect of taxa *j* on taxa *i,* while *ϵ_ik_* denotes the effect of antimicrobial *k* on taxon *i* and *E_k_* is a binary variable denoting whether antibiotic *k* was administered at time *t.*

The signs of *α_ij_* and *α_ij_* together define the nature of the interaction between each pair of taxa *i* and *j*. Specifically, when both terms are positive (+/+) the interaction is mutualistic and when both are negative (−/−) the interaction is competitive. When one taxa gains a benefit at the expense of the other (+/−) we term the interaction exploitative, when one taxa benefits while the other is unaffected the interaction is commensal (+/0), and when one taxa is harmed while the other is unaffected the interaction is amensal (−/0).

This gLV model can further be discretized and linearized such that it is equivalent to,

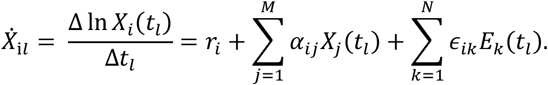

Where *X_j_(t_l_)* now represents the geometric mean of taxon *j* across the time window *l*. In this linear form, we can now fit each of the parameters *r*, *α*, and *ϵ* using simple linear regression.

To avoid overfitting, and ensure we focused on those interactions that play the most significant role in shaping community dynamics we used a conservative form of variable selection. Specifically, we implemented a Bayesian spike-and-slab regression procedure, which forced the majority of our interaction parameters to zero. We constructed this by setting the prior for each interaction parameter to have a point mass at zero combined with a half-Cauchy random variable. We allowed the microbe-microbe and drug-microbe interactions to regularize separately. In practice, our regression model was thus defined as,

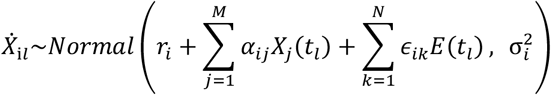

Dropping the subscript *i* for ease of reading, our spike-and-slab regularization procedure for each taxon, *i,* was implemented using the following hierarchical priors,

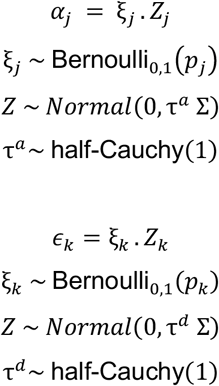

Where Σ was a Zellner prior, 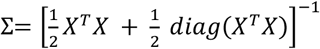, adapted to ensure invertibility. Notably though, we inferred similar results instead simply using the identity matrix.

Following Bucci et al, to enforce non-negative growth rates, we assumed a truncated Normal prior on the intrinsic growth rate of each taxon,

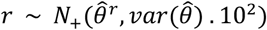

with the hyperparameters calculated from the pseudoinverse solution to the linear regression 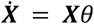, ie by solving 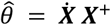. Finally, again following Bucci et al, we allowed a taxon-specific variance, 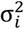, set simply as an inflated half-normal distribution.

To reduce the complexity of our model, before fitting we filtered our data to only include those taxa that reached 1% of the total population in at least 10% of samples. We also removed any drugs that were administered on fewer than five days across the entire 13 infant cohort. To ensure we did not bias our fits against low abundance taxa, we normalized our independent variables to each have zero mean and standard deviation of one. We then solved the above model using the Python package *pymc3*, drawing 10,000 samples after a burn in of 2500. Finally, we further filtered our results by discounting any normalized parameters whose magnitude fell below 0.01.

### Section 4 – Validation of microbe-microbe interactions

#### Strain isolation

We used selective agar media to isolate microbes of targeted taxa. To isolate *Candida*, we used Yeast Extract–Peptone–Dextrose (YPD) supplemented with antibacterial cocktail (64 ug/mL aztreonam and 5 ug/mL vancomycin final concentration). To isolate *Klebsiella* and *Escherichia*, we used MacConkey supplemented with antifungal cocktail (0.05 ug/mL micafungin and 50ug/mL cycloheximide final concentration). To isolate *Staphylococcus* and *Enterococcus*, we used Brain Heart Infusion (BHI) supplemented with aztreonam and antifungal cocktail. We selected stool samples from 5 NICU subjects (NICU_073, 091, 175, 203 and 260), homogenized samples in sterile PBS and plated out dilutions on each of these media. After 1~2 day incubation aerobic at 37C, colonies of representative morphologies were selected for full-length 16S PCR amplification and Sanger sequencing. The full-length 16S sequences were blasted to identify species-level taxonomies and the chromatogram data were used to map back to the Miseq OTU sequences (Supplemental Table 13). Non-redundant strains were saved in glycerol stocks and recovered on BHI agar media before use.

#### *In vitro* co-culture validation

To facilitate co-culture of taxonomically diverse bacteria and fungi in a nutrient tractable condition, we developed a common defined medium YNBGS based on Yeast Nitrogen Base (YNB): 5.0 g/L glucose, 2.0 g/L Casamino acids, 0.1 g/L L-tryptophan, 20 mg/L adenine and 20 mg/L guanine were supplemented to 6.7 g/L YNB (Millipore Sigma, Cat Y1250), suspended in water and filter sterilized. All co-culture experiments were done in the YNBGS broth at 37C static, under aerobic conditions for 24 hours unless otherwise stated (we used aerobic conditions as the neonatal gut is known to be relatively aerobic). Recovered bacteria and fungi were suspended directly in YNBGS and diluted to OD 10^−3, before mixing with each other. Note that because *Candida* replicated slower and cell counts per OD were lower than the bacteria, when testing the effect of *Candida* on others we first cultured *Candida* in YNB starting from OD 0.1 at 37C for 24 hours (final OD 0.5~0.7), before mixing 1:1 (volume ratio) with bacterial suspension. We spotted 2.5 uL of 10-fold serial dilutions of the cultures, either inoculation inputs or mono-/co-culture outputs, onto the aforementioned selective media, and used the CFU counts to quantify growth fitness effects. Each *in vitro* result shown (Fig. 3d, f, Supplemental Figure 9, Supplemental Table 14) was representative of at least two independent experiments.

#### *In vivo* co-colonization validation

We used specific pathogen free (SPF) 6~8 weeks old female mice (Jackson Laboratory, Bar Harbor, Maine) for *in vivo* validation under the animal protocol 18-02-3637R, approved by the Institutional Animal Care and Use Committee at Boston Children’s Hospital. Five mice per group were co-housed and tail marked before colonization. To facilitate colonization of exogenous bacteria and fungi of interest in the mouse gut, we applied a 3-day antibiotics treatment (vancoymin+metronidazole+ampicillin cocktail, 1 g/L each in drinking water) prior to the 1st oral gavage, and maintained ampicillin drinking water (replaced fresh every 3 days) afterwards during the whole experiment. In Fig. 3, we used *K. pneumoniae*, *S. epidermidis* and *C. albicans* strains all isolated from infant NICU_203 and *C. parapsilosis* from infant NICU_091 (all clones were confirmed to be resistant to at least 100 ug/mL ampicillin). We cultured bacteria and fungi in BHI broth shaking overnight at 37C, concentrated in 200 uL sterile PBS for oral gavage into each mouse. Note that a lower inoculation was used for *K. pneumoniae* than others as this bacterium more readily and rapidly established colonization in the mouse gut; and for *S. epidermidis* pre-treatment colonization only, we performed a reinforcement gavage of *S. epidermidis*, as we noticed inconsistent levels of *S. epidermidis* in the stool samples 1-day after the 1st gavage (Supplemental Table 15). Daily stool samples before, at and after gavage were collected for each mouse (a 12-hr time point was also collected for Fig. 3e). To evaluate the colonization level of targeted strains, collected stool samples were suspended in sterile PBS (100 uL per 1 mg stool) by vortexing for > 5 min, and 10-fold serial dilutions were plated onto the aforementioned selective media for CFU counting. We corroborated our CFU data by using MK-SpikeSeq on the collected stool samples, focusing on taxa of the gavaged bacteria and fungi (Supplemental Figure 10, Supplemental Table 15).

## Acknowledgements

We thank all of the infants and their families who participated in the study. We also thank Joao Xavier, Jose Ordovas-Montanes, Olivier Cunrath, and members of the Rakoff-Nahoum Lab for helpful discussions, and Linnea Martin for assistance with sample collection. K.Z.C. is funded by a Sir Henry Wellcome Postdoctoral Research Fellowship (grant 201341/A/16/Z) and a University of Manchester Presidential Fellowship. S.R-N. is supported by a Career Award for Medical Scientists from the Burroughs Wellcome Fund, a Pew Biomedical Scholarship, a Basil O’Connor Starter Scholar Award from the March of Dimes, K08AI130392-01 and a NIH Director’s New Innovator Award DP2GM136652.

## References

1. Charbonneau, M. R. et al. Human developmental biology viewed from a microbial perspective. Nature 535, 48–55 (2016).

2. Bäckhed, F. et al. Dynamics and stabilization of the human gut microbiome during the first year of life. Cell Host Microbe 17, 690–703 (2015).

3. Stewart, C. J. et al. Temporal development of the gut microbiome in early childhood from the TEDDY study. Nature 562, 583–588 (2018).

4. Derrien, M., Alvarez, A. S. & de Vos, W. M. The Gut Microbiota in the First Decade of Life. Trends in Microbiology 27, 997–1010 (2019).

5. Yatsunenko, T. et al. Human gut microbiome viewed across age and geography. Nature 486, 222–227 (2012).

6. Lim, E. S. et al. Early life dynamics of the human gut virome and bacterial microbiome in infants. Nat. Med. 21, 1228–1234 (2015).

7. Palmer, C., Bik, E. M., DiGiulio, D. B., Relman, D. A. & Brown, P. O. Development of the human infant intestinal microbiota. PLoS Biol. 5, 1556–1573 (2007).

8. Lynch, S. V. & Pedersen, O. The Human Intestinal Microbiome in Health and Disease. N. Engl. J. Med. 375, 2369–2379 (2016).

9. Honda, K. & Littman, D. R. The microbiota in adaptive immune homeostasis and disease. Nature 535, 75–84 (2016).

10. Fischbach, M. A. Microbiome: Focus on Causation and Mechanism. Cell 174, 785–790 (2018).

11. Widder, S. et al. Challenges in microbial ecology: Building predictive understanding of community function and dynamics. ISME Journal 10, 2557–2568 (2016).

12. Vrancken, G., Gregory, A. C., Huys, G. R. B., Faust, K. & Raes, J. Synthetic ecology of the human gut microbiota. Nature Reviews Microbiology 17, 754–763 (2019).

13. Walter, J., Armet, A. M., Finlay, B. B. & Shanahan, F. Establishing or Exaggerating Causality for the Gut Microbiome: Lessons from Human Microbiota-Associated Rodents. Cell 180, 221–232 (2020).

14. Wolfe, B. E., Button, J. E., Santarelli, M. & Dutton, R. J. Cheese rind communities provide tractable systems for in situ and in vitro studies of microbial diversity. Cell 158, 422–433 (2014).

15. Connell, J. H. & Slatyer, R. O. Mechanisms of Succession in Natural Communities and Their Role in Community Stability and Organization. The American Naturalist 111, 1119–1144

16. Bertness, M. D. & Callaway, R. Positive interactions in communities. Trends in Ecology and Evolution 9, 191–193 (1994).

17. Shade, A. et al. Macroecology to Unite All Life, Large and Small. Trends in Ecology and Evolution 33, 731–744 (2018).

18. Gregory, K. E. et al. Influence of maternal breast milk ingestion on acquisition of the intestinal microbiome in preterm infants. Microbiome 4, 68 (2016).

19. Gibson, M. K. et al. Developmental dynamics of the preterm infant gut microbiota and antibiotic resistome. Nat. Microbiol. 1, 16024 (2016).

20. Dibartolomeo, M. E. & Claud, E. C. The Developing Microbiome of the Preterm Infant. Clinical Therapeutics 38, 733–739 (2016).

21. La Rosa, P. S. et al. Patterned progression of bacterial populations in the premature infant gut. Proc. Natl. Acad. Sci. 111, 12522–12527 (2014).

22. Costello, E. K., Carlisle, E. M., Bik, E. M., Morowitz, M. J. & Relman, D. A. Microbiome assembly across multiple body sites in low-birthweight infants. MBio 4, e00782–13 (2013).

23. Stewart, C. J. et al. Temporal bacterial and metabolic development of the preterm gut reveals specific signatures in health and disease. Microbiome 4, 67 (2016).

24. Pammi, M. et al. Intestinal dysbiosis in preterm infants preceding necrotizing enterocolitis: A systematic review and meta-analysis. Microbiome 5, (2017).

25. Gasparrini, A. J. et al. Persistent metagenomic signatures of early-life hospitalization and antibiotic treatment in the infant gut microbiota and resistome. Nat. Microbiol. 4, 2285–2297 (2019).

26. Reynolds, L. A. & Finlay, B. B. Early life factors that affect allergy development. Nature Reviews Immunology 17, 518–528 (2017).

27. Gensollen, T., Iyer, S. S., Kasper, D. L. & Blumberg, R. S. How colonization by microbiota in early life shapes the immune system. Science 352, 539–544 (2016).

28. Pasolli, E. et al. Extensive Unexplored Human Microbiome Diversity Revealed by Over 150,000 Genomes from Metagenomes Spanning Age, Geography, and Lifestyle. Cell 176, 649–662.e20 (2019).

29. Proctor, L. M. et al. The Integrative Human Microbiome Project. Nature 569, 641–648 (2019).

30. Nash, A. K. et al. The gut mycobiome of the Human Microbiome Project healthy cohort. Microbiome 5, 153 (2017).

31. Limon, J. J., Skalski, J. H. & Underhill, D. M. Commensal Fungi in Health and Disease. Cell Host and Microbe 22, 156–165 (2017).

32. Koskinen, K. et al. First insights into the diverse human archaeome: Specific detection of Archaea in the gastrointestinal tract, lung, and nose and on skin. MBio 8, 1–17 (2017).

33. Durán, P. et al. Microbial Interkingdom Interactions in Roots Promote Arabidopsis Survival. Cell 175, 973–983.e14 (2018).

34. Carr, A., Diener, C., Baliga, N. S. & Gibbons, S. M. Use and abuse of correlation analyses in microbial ecology. ISME J. 1 (2019). doi:10.1038/s41396-019-0459-z

35. Contijoch, E. J. et al. Gut microbiota density influences host physiology and is shaped by host and microbial factors. Elife 8, (2019).

36. Vandeputte, D. et al. Quantitative microbiome profiling links gut community variation to microbial load. Nature 551, 507–511 (2017).

37. Stämmler, F. et al. Adjusting microbiome profiles for differences in microbial load by spike-in bacteria. Microbiome 4, 28 (2016).

38. Buffie, C. G. et al. Precision microbiome reconstitution restores bile acid mediated resistance to Clostridium difficile. Nature 517, 205–208 (2015).

39. Kamra, A. et al. Exfoliated Colonic Epithelial Cells: Surrogate Targets for Evaluation of Bioactive Food Components in Cancer Prevention. J. Nutr. 135, 2719–2722 (2005).

40. Ishwaran, H. & Rao, J. S. SPIKE AND SLAB VARIABLE SELECTION: FREQUENTIST AND BAYESIAN STRATEGIES. Ann. Stat. 33, 730–773 (2005).

41. Gonze, D., Coyte, K. Z., Lahti, L. & Faust, K. Microbial communities as dynamical systems. Curr. Opin. Microbiol. 44, (2018).

42. Bucci, V. et al. MDSINE: Microbial Dynamical Systems INference Engine for microbiome time-series analyses. Genome Biol. 17, 121 (2016).

43. Freilich, M. A., Wieters, E., Broitman, B. R., Marquet, P. A. & Navarrete, S. A. Species co-occurrence networks: Can they reveal trophic and non-trophic interactions in ecological communities? Ecology 99, 690–699 (2018).

44. Friedman, J. & Alm, E. J. Inferring Correlation Networks from Genomic Survey Data. PLoS Comput. Biol. 8, e1002687 (2012).

45. Faust, K. et al. Microbial Co-occurrence Relationships in the Human Microbiome. PLoS Comput. Biol. 8, e1002606 (2012).

46. Watts, S. C., Ritchie, S. C., Inouye, M. & Holt, K. E. FastSpar: rapid and scalable correlation estimation for compositional data. Bioinformatics 35, 1064–1066 (2019).

