## Supplemental figures and tables for "Multi-kingdom quantitation reveals distinct ecological drivers of predictable early-life microbiome assembly"

### Section 5 – Supplementary Figures and Tables

Figures

Supplemental Figure 1. MK-SpikeSeq outperforms other quantitation methods in cross-kingdom sensitivity and specificity

Supplemental Figure 2. MK-SpikeSeq captures key ecological dynamics in mock communities

Supplemental Figure 3. Bacterial samples cluster based on composition and infant age, but not diet, delivery mode, or gender

Supplemental Figure 4. Fungal community composition does not map to infant age, diet, gender or delivery mode

Supplemental Figure 5. Bacterial community dynamics exhibit more temporal structure than fungal communities

Supplemental Figure 6. Trends in total microbial loads for all three kingdoms

Supplemental Figure 7. Microbe-microbe interactions are predominantly asymmetric, while antimicrobials primarily inhibit their target kingdom

Supplemental Figure 8. Inferring interactions from relative abundance data generates misleading results

Supplemental Figure 9. rDNA-based measurement of *in vivo* colonization using MK-SpikeSeq

Tables

Supplemental Table 1. Primers used in the study

Supplemental Table 2. Absence of select spike-in taxa in public data depositories.

Supplemental Table 3. Comparison of absolute quantification methods using test samples.

Supplemental Table 4. Sensitivity of MK-SpikeSeq versus qPCR.

Supplemental Table 5. Robustness of MK-SpikeSeq versus qPCR.

Supplemental Table 6. Testing accuracy of MK-SpikeSeq in capturing community changes.

Supplemental Table 7. NICU cohort stool samples sequenced in the study.

Supplemental Table 8. Bacteria OTU table in the 1st phase.

Supplemental Table 8. Bacteria OTU table in the 1st phase.

Supplemental Table 9. Archaea OTU table in the 1st phase.

Supplemental Table 10. Fungi OTU table in the 1st phase.

Supplemental Table 11. Bacteria OTU table in the 2nd phase.

Supplemental Table 12. Fungi OTU table in the 2nd phase.

Supplemental Table 13. Strains isolated from select NICU subjects.

Supplemental Table 14. CFU fold growth in *in vitro* co-culture experiments.

Supplemental Table 15. CFU- and rDNA-based quantification of *in vivo* co-colonizations.

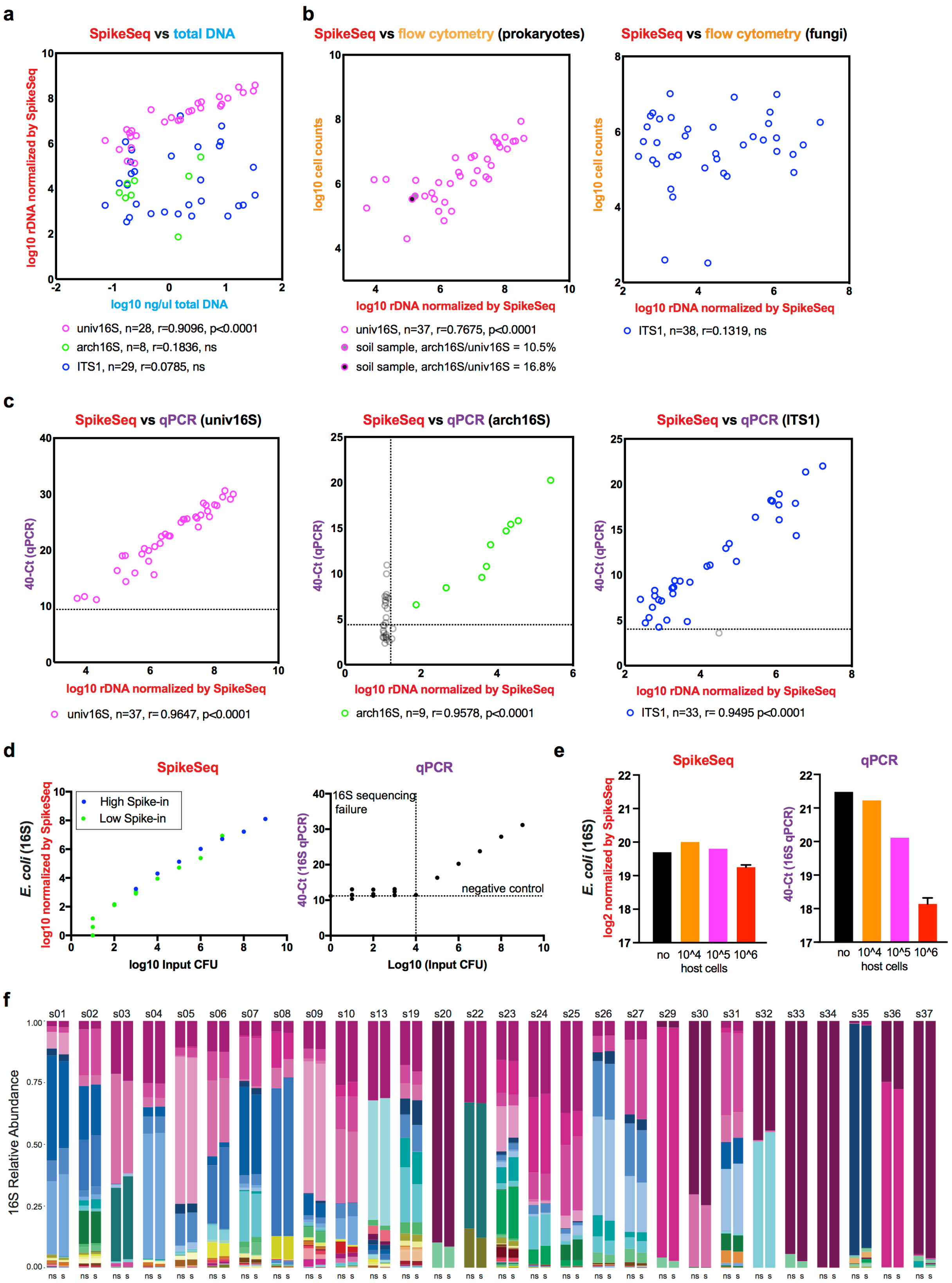
a
SpikeSeq vs total DNA
log10 rDNA normalized by SpikeSeq
log10 ng/ul total DNA
univ16S, n=28, r=0.9096, p<0.0001
arch16S, n=8, r=0.1836, ns
ITS1, n=29, r=0.0785, ns
b
SpikeSeq vs flow cytometry (prokaryotes)
log10 cell counts
log10 rDNA normalized by SpikeSeq
univ16S, n=37, r=0.7675, p<0.0001
soil sample, arch16S/univ16S = 10.5%
soil sample, arch16S/univ16S = 16.8%
SpikeSeq vs flow cytometry (fungi)
log10 cell counts
log10 rDNA normalized by SpikeSeq
ITS1, n=38, r=0.1319, ns
c
SpikeSeq vs qPCR (univ16S)
40-Ct (qPCR)
log10 rDNA normalized by SpikeSeq
univ16S, n=37, r= 0.9647, p<0.0001
SpikeSeq vs qPCR (arch16S)
40-Ct (qPCR)
log10 rDNA normalized by SpikeSeq
arch16S, n=9, r= 0.9578, p<0.0001
SpikeSeq vs qPCR (ITS1)
40-Ct (qPCR)
log10 rDNA normalized by SpikeSeq
ITS1, n=33, r= 0.9495 p<0.0001
d
SpikeSeq
E. coli (16S) log10 normalized by SpikeSeq
High Spike-in
Low Spike-in
log10 Input CFU
qPCR
40-Ct (16S qPCR)
16S sequencing failure
negative control
Log10 (Input CFU)
e
SpikeSeq
E. coli (16S) log2 normalized by SpikeSeq
no
10^4
10^5
10^6
host cells
qPCR
40-Ct (16S qPCR)
no
10^4
10^5
10^6
host cells
f
s01 s02 s03 s04 s05 s06 s07 s08 s09 s10 s13 s19 s20 s22 s23 s24 s25 s26 s27 s29 s30 s31 s32 s33 s34 s35 s36 s37
16S Relative Abundance
ns s

**Figure S1. MK-SpikeSeq outperforms other quantitation methods in cross-kingdom sensitivity and specificity. a**, The kingdom-specific abundances of test samples measured by MK-SpikeSeq are compared with the total DNA concentrations measured by the PicoGreen assay. Pearson correlation tests show that total DNA yields mostly only reflect bacterial community abundances. **b**, The flow cytometry cell enumerations using gating strategies targeted for either prokaryotes or fungi are compared with MK-SpikeSeq quantifications. For prokaryotic enumerations, two soil samples are highlighted due to their high archaeal loads that cannot be distinguished from bacterial counts by flow cytometry. For fungi enumerations, shown are results using one gating strategy; attempts using two additional gating strategies show similar over-estimation of fungal counts (Supplemental Table 3). ns, not significant. **c**, The kingdom-specific qPCR Ct values are compared with MK-SpikeSeq quantifications. Horizontal dashed lines show the limit of detection using qPCR, based on the negative control (DNA extraction of water); vertical dashed line shows the limit of detection using MK-SpikeSeq, based on the normalization of minimal one non-spike-in arch16S read against the average arch16S sequencing depth. Samples below either limit of detection are excluded from correlational analyses. **d**, Comparison of sensitivity between SpikeSeq and qPCR using 10-fold serial dilutions of *E. coli*. SpikeSeq showed 100~1000-fold increased sensitivity over qPCR in low microbial abundance samples (detecting as few as 10 bacterial cells). For SpikeSeq, two levels of spike-in were used to cover the whole range of detection under the sequencing depth of 10~100k reads per sample (see Supplemental Information). For qPCR, horizontal dashed line shows Ct value of the negative control and vertical dashed line shows the threshold below which pooled 16S sequencing yielded less than 100 reads per sample (sequencing failed likely due to too low signal). **e**, Comparison of robustness to host cell background between SpikeSeq and qPCR using test samples with fixed *E. coli* abundance and variable number of Caco-2 colonic cells. SpikeSeq detected consistent microbial abundances in samples with high host-cell background that were under-measured by 10-fold using qPCR. **f**, Comparison of 16S genus-level profiles sequenced with (s) or without (ns) spike-in shows largely unaltered community compositions having exogenous spike-in.

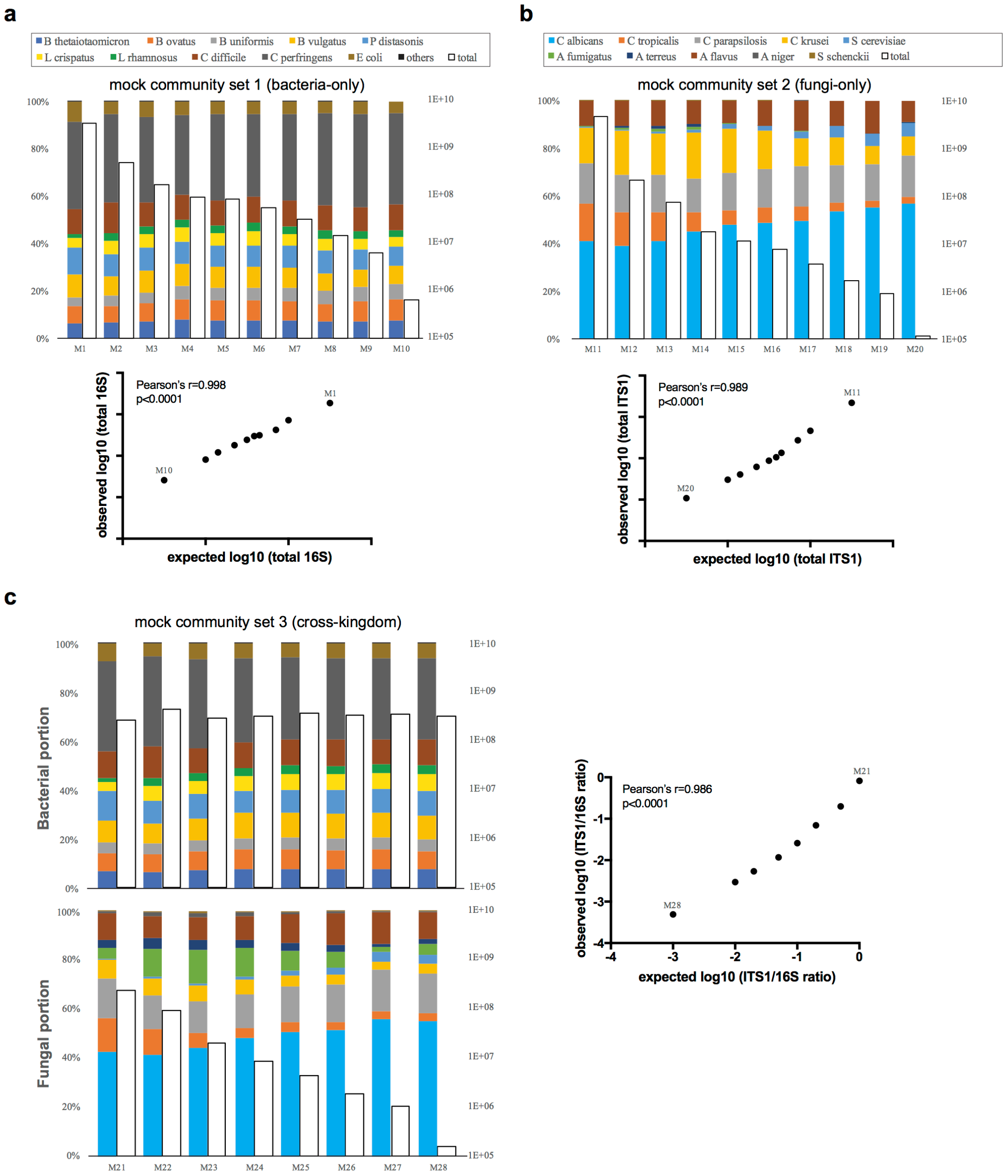


**Figure S2. MK-SpikeSeq captures key ecological dynamics in mock communities. a, b**, A set of single-kingdom mock communities with a fixed composition of 10 bacterial or 10 fungal species and variable total microbial loads, characterized by relative composition (colored bars) and total abundance (empty bars) measured by SpikeSeq. Correlations between expected (based on known dilution factors) and SpikeSeq-measured total abundances show that SpikeSeq reliably detects intra-kingdom absolute abundance. **c**, A set of dual-kingdom communities with different ratios of the defined bacterial and fungal sub-communities, characterized by MK-SpikeSeq. Correlations between expected (based on known dilution factors) and MK-SpikeSeq-measured bacteria-fungi ratios show that MK-SpikeSeq reliably captures cross-kingdom absolute abundance.

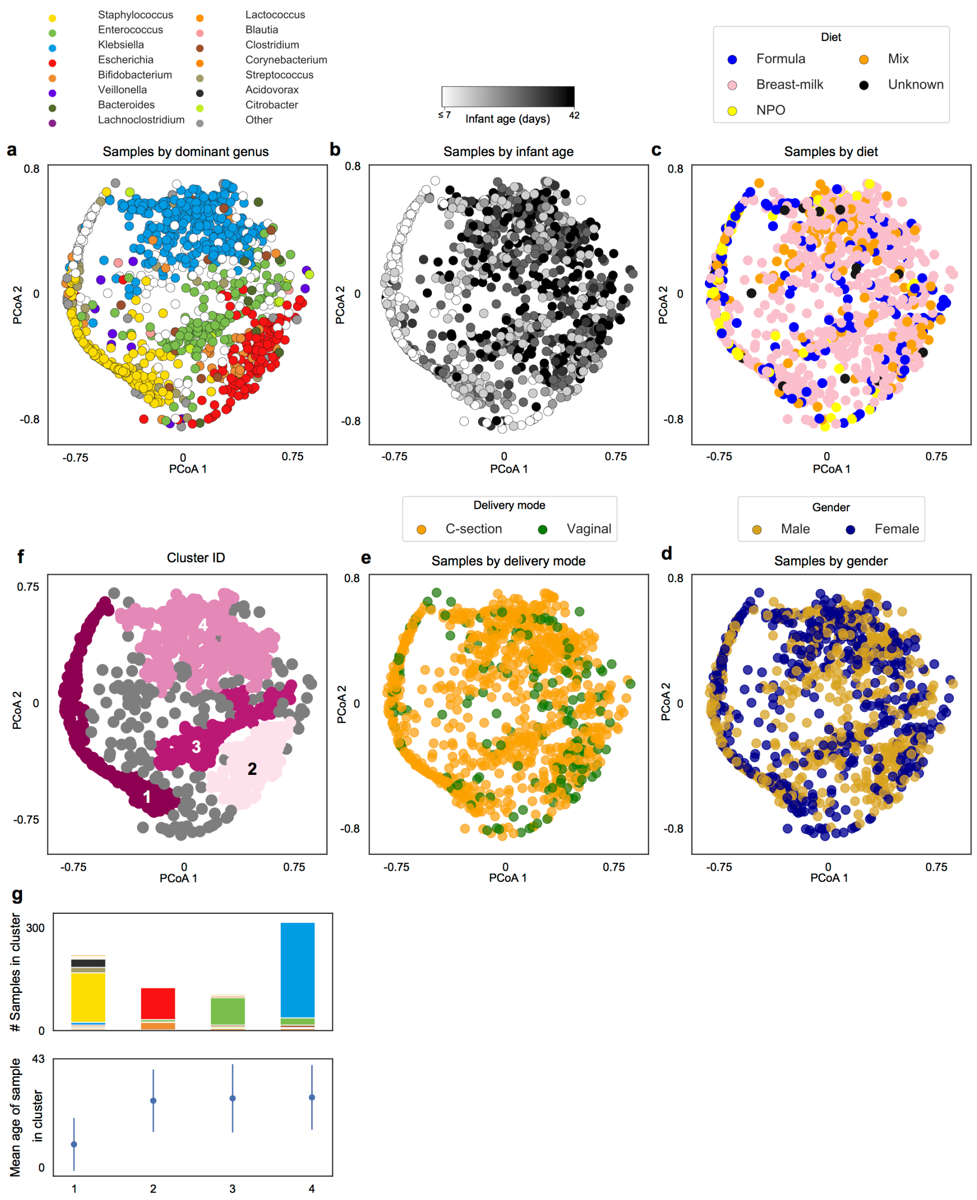


**Figure S3. Bacterial samples cluster based on composition and infant age, but not diet, delivery mode, or gender. a,** Principle Coordinate Analysis (PCoA) based on Bray-Curtis dissimilarities of bacterial community composition between samples (genus level). Samples colored by dominant taxa or white when diversity is high (IS>4). **b,** PCoA colored by infant age. **c,** PCoA colored by infant diet close. **d,** PCoA with samples colored by infant gender. e, PCoA with samples colored by delivery mode. **f,** PCoA with samples colored by cluster membership, calculated using DBSCAN algorithm. **g,** Stacked bars represent distribution of dominant genus within each cluster and dot plots illustrate average day of life of samples within each cluster. Kruskal-Wallis test with Bonferroni correction showed statistically significant differences in day of life of samples between clusters (Chi square = 254, p-value << 0.05, df = 3), error bars show standard deviation.

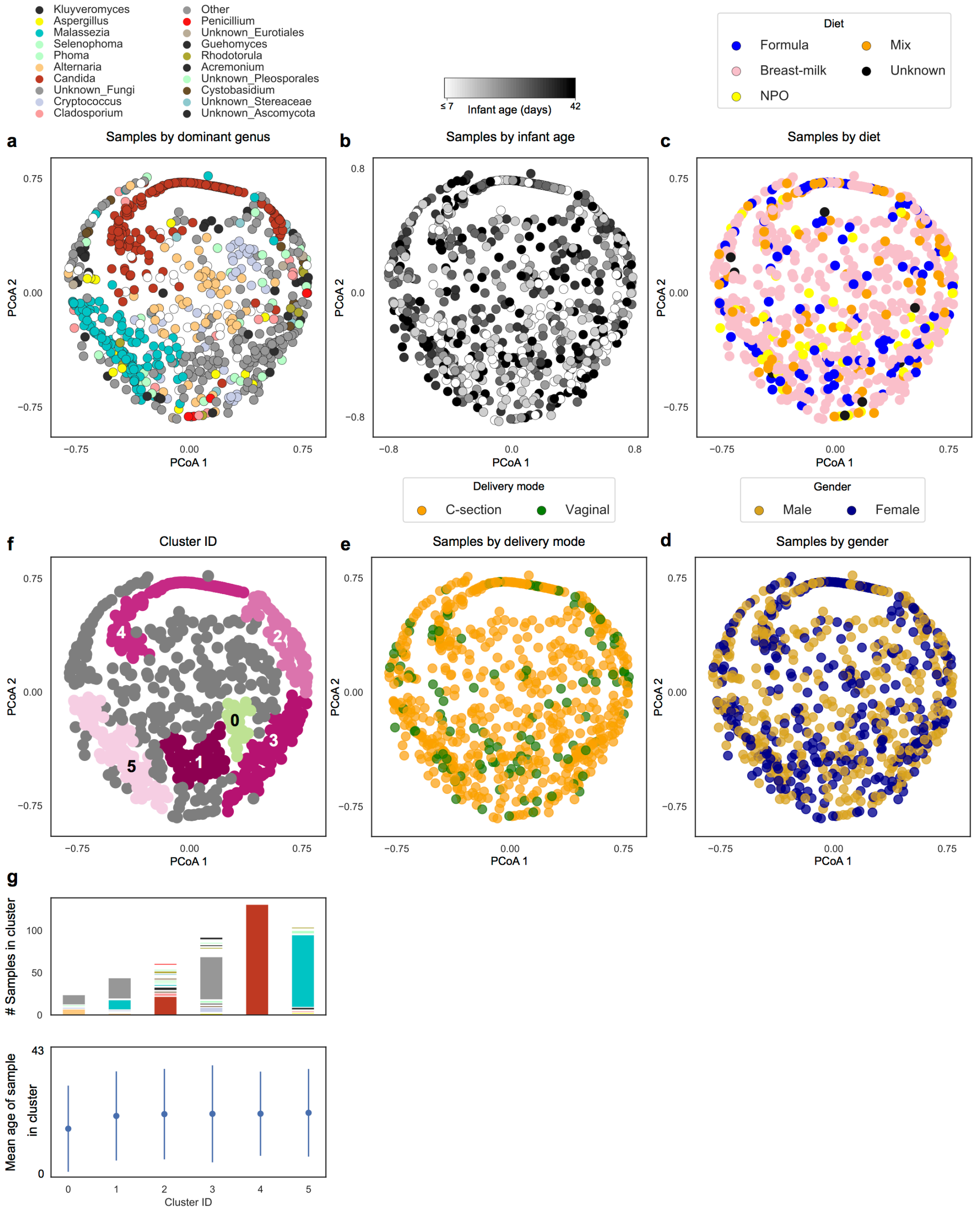


**Figure S4. Fungal community composition does not map to infant age, diet, gender or delivery mode. a,** Principle Coordinate Analysis (PCoA) based on Bray-Curtis dissimilarities of fungal community composition between samples (genus level). Samples colored by dominant taxa or white when diversity is high (IS>4). **b,** PCoA colored by infant age. **c,** PCoA colored by infant diet close. **d,** PCoA with samples colored by infant gender. **e,** PCoA with samples colored by delivery mode. **f,** PCoA with samples colored by cluster membership, calculated using DBSCAN algorithm. **g,** Stacked bars represent distribution of dominant genus within each cluster and dot plots illustrate average day of life of samples within each cluster. Kruskal-Wallis test with Bonferroni correction showed no statistically significant differences in day of life of samples between clusters (Chi square = 3.06, p-value = 0.69, df = 5), error bars show standard deviation.

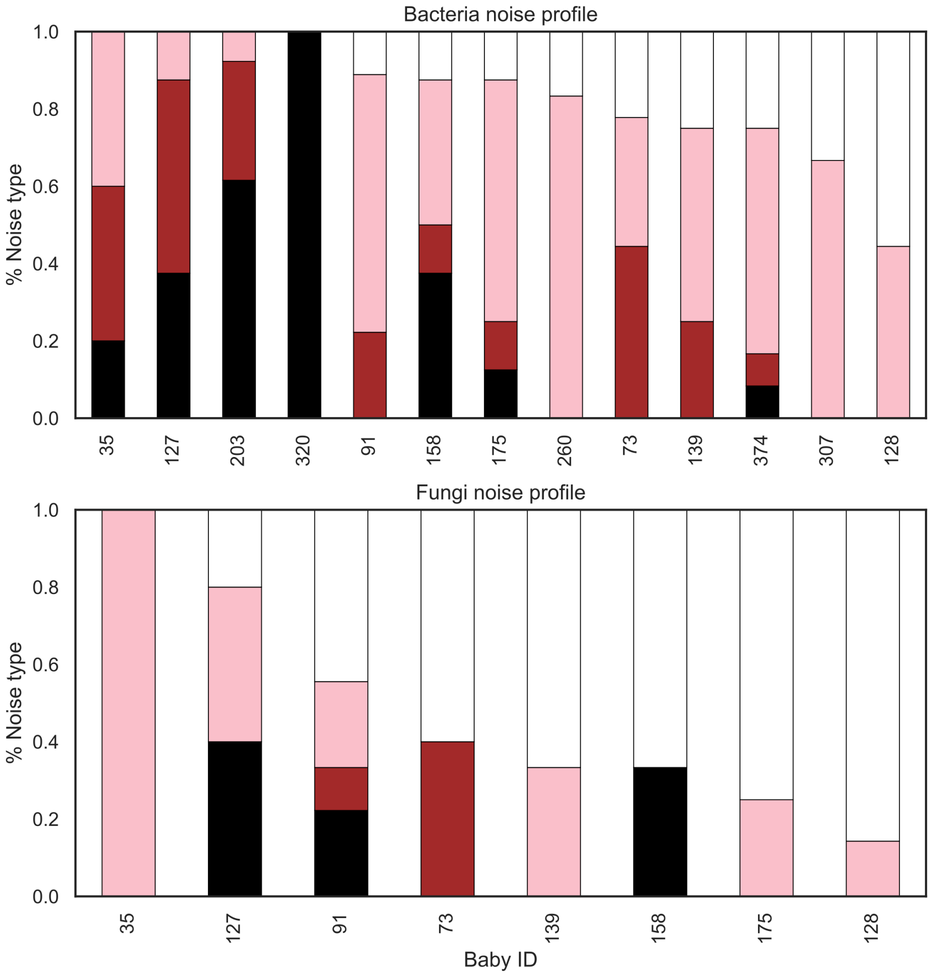


**Figure S5. Bacterial community dynamics exhibit more temporal structure than fungal communities.** Stacked bars indicate the proportion on genera exhibiting each noise type per infant. Dark noise indicates increasing temporal dependence, with white noise suggesting temporal dynamics are entirely random. Infant bacterial communities show a relatively high proportion of temporal structure, with on average only x% of genera exhibiting white noise. By contrast, fungal dynamics exhibited far less temporal structure, with on average y% of genera displaying white noise, and thus no time dependent dynamics. Notably, 5 infants mycobiomes were so chaotic they could not be classified.

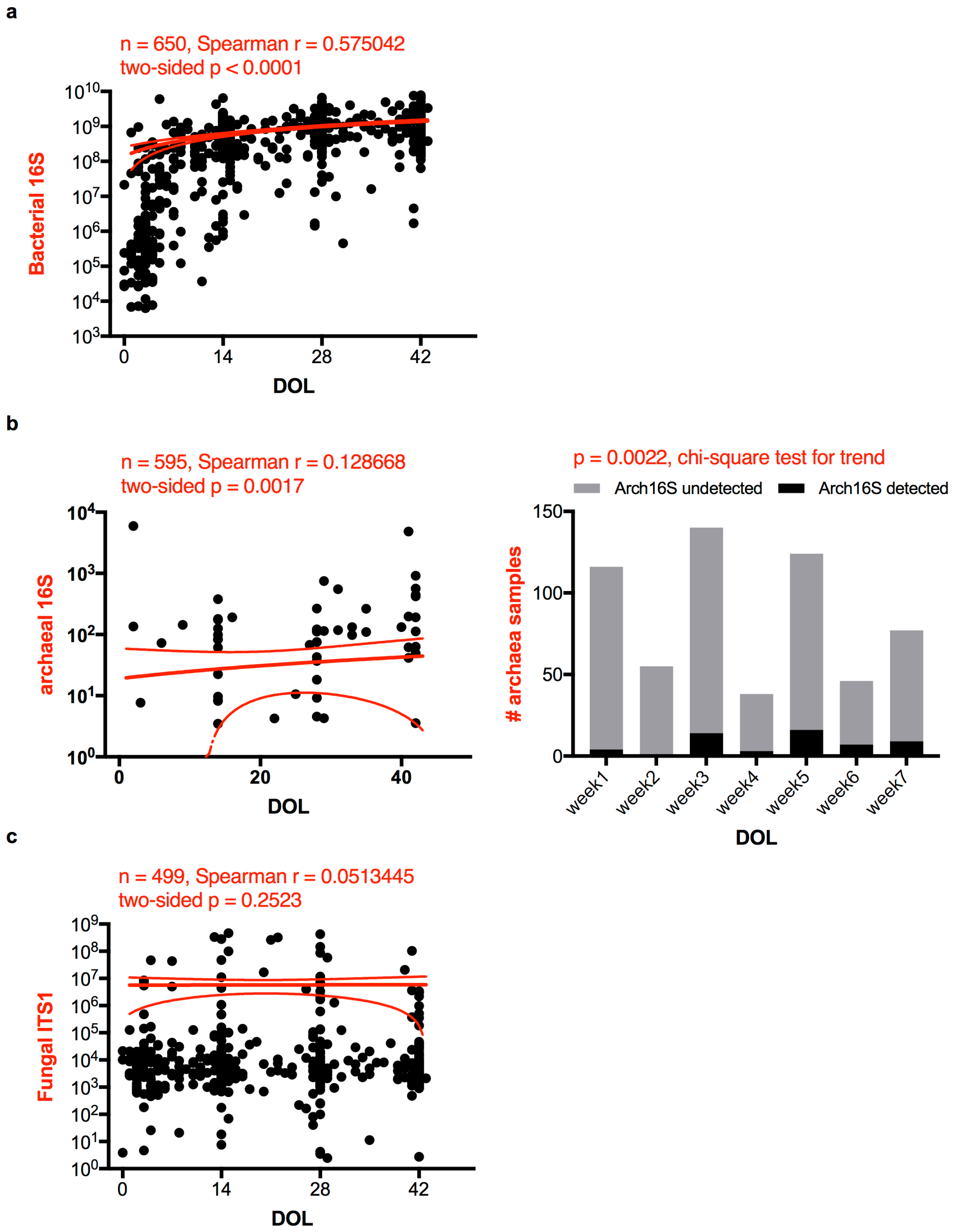


**Figure S6. Trends in total microbial loads for all three kingdoms.** Scatter plots of rDNA-based total abundances of individual kingdoms against infant day of life (DOL), measured by MK-SpikeSeq in the first phase Nextseq sequencing. The red lines denote the linear regression fit and the 90% confidence bands of the best-fit line of each dataset. Spearman correlations show that bacterial and archaeal, but not fungal, loads are positively associated with infant age. Samples with undetectable kingdom-specific rDNA signal are not plotted. For archaea that show scarce signal in the cohort (**b**), a separate presence/absence plot and chi-square test of binned samples also show a positive correlation between archaeal loads and infant age.

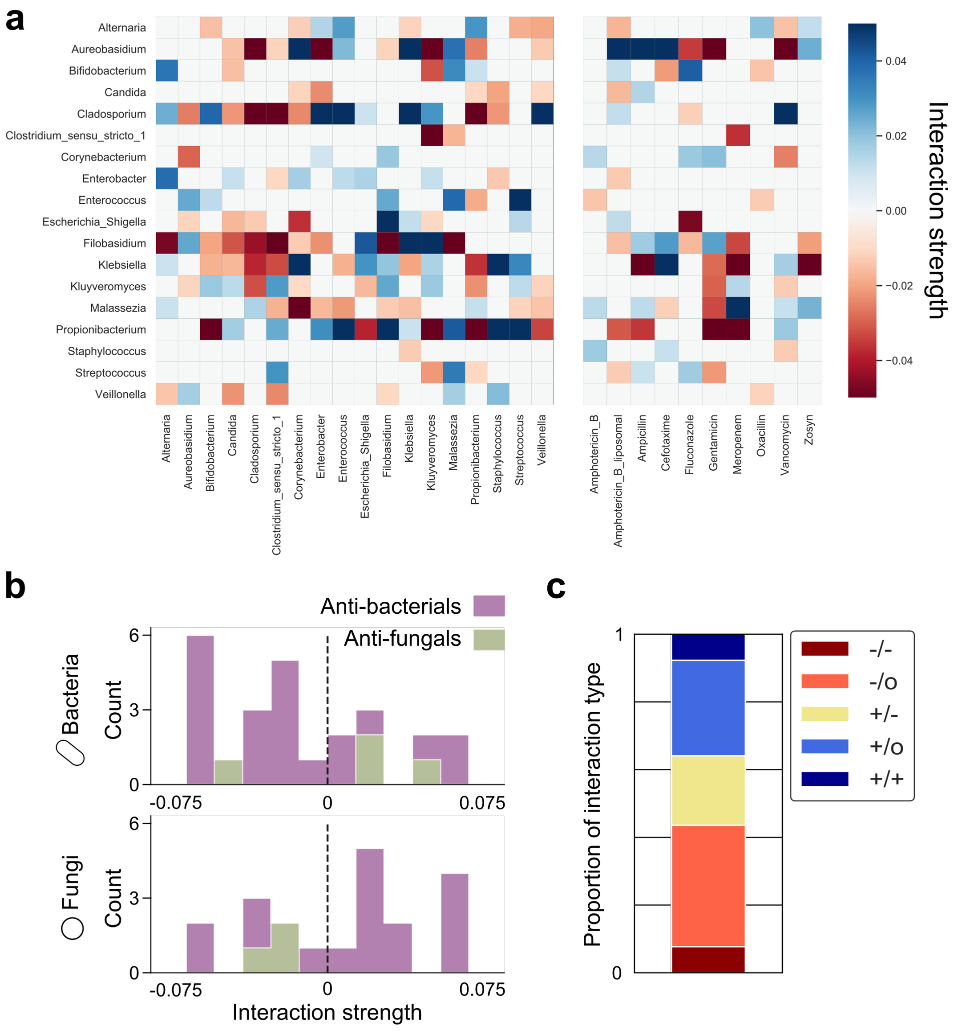


**Figure S7. Microbe-microbe interactions are predominantly asymmetric, while antimicrobials primarily inhibit their target kingdom**. **a,** Heatmap plotting interactions inferred by the gLV model. Each row of the heatmap illustrates the effect upon the target genera by other members of the gut community (left columns) or documented usage of antimicrobials according the clinical metadata (right columns). **b,** Histogram of individual antibacterial (purple) or antifungal (green) interaction strengths, split by kingdom. As expected, antibacterials primarily inhibit bacteria, and antifungals primarily inhibit fungi. **c,** Stacked bar illustrates the proportion of different interaction types occurring between genera. Over 80% of interactions are asymmetric, being either exploitative (+/-), commensal (+/0), or amensal (-/0).

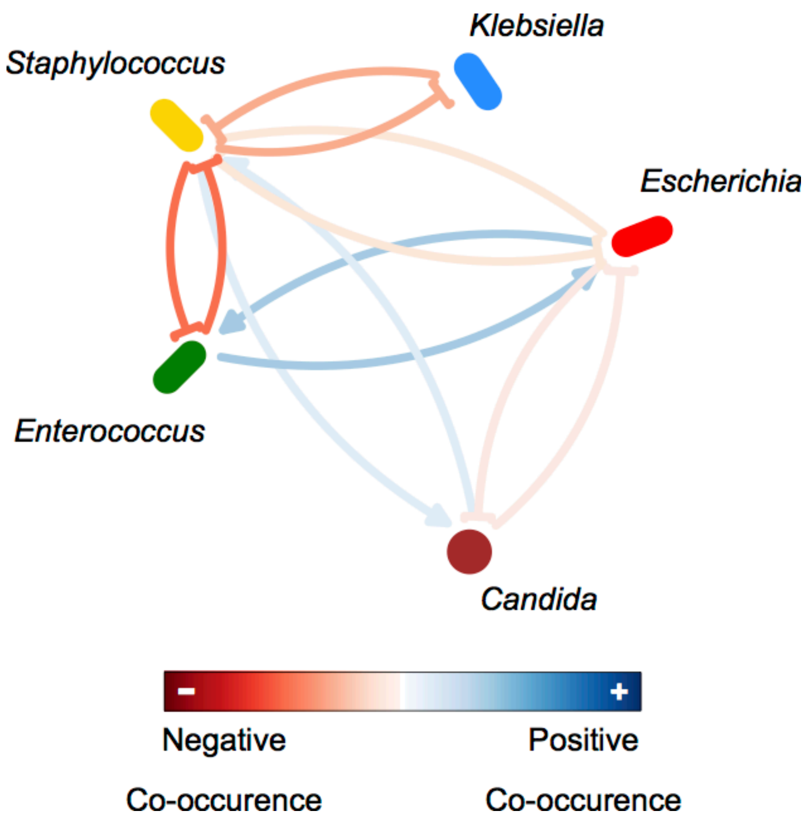


**Figure S8. Inferring interactions from relative abundance data generates misleading results**. To confirm the value of our absolute abundance methods, we inferred inter-genus interactions from relative abundance data alone using the FastSpar[48] algorithm. This approach robustly identifies co-occurrence relationships between different microbial taxa in a manner that accounts for the compositional nature of relative abundance data. Notably, correlation networks cannot infer asymmetric interactions thus this approach cannot detect the exploitation of *Staphylococcus* by *Klebsiella*. It also erroneously infers that *Staphylococcus* increases the growth of *Candida*, and cannot detect the inhibition of *Klebsiella* by *Candida* or *Enterococcus*.

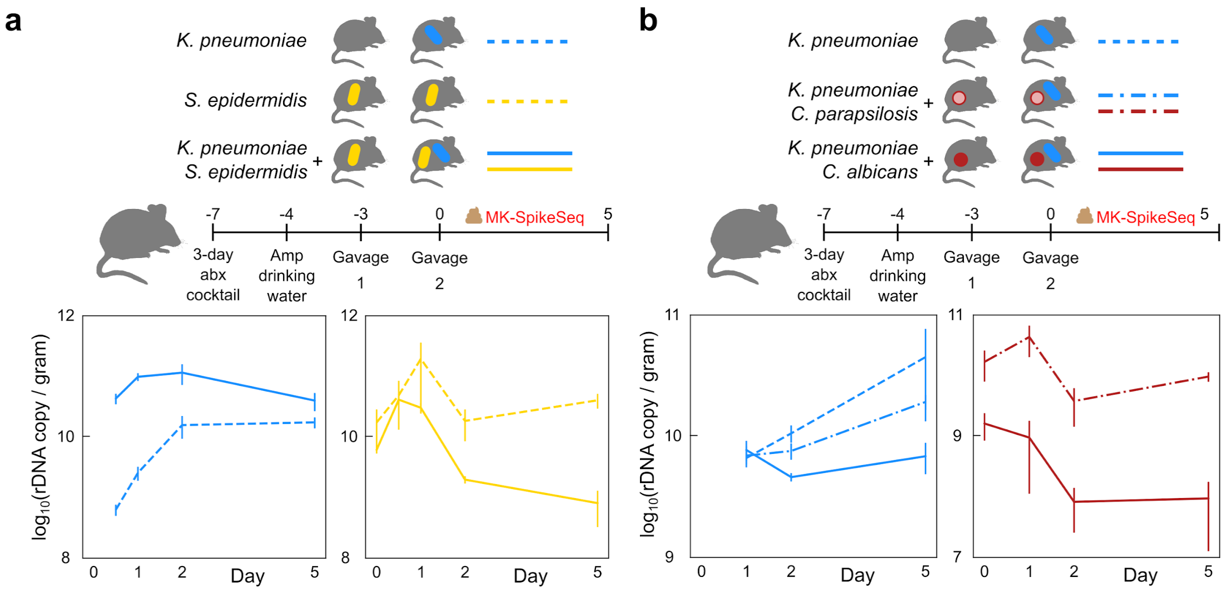


**Figure S9. rDNA-based measurement of *in vivo* colonization using MK-SpikeSeq.** Biological replicate samples of mouse stools characterized by CFU counting of strains of interest were subjected to MK-SpikeSeq to determine rDNA-based absolute abundances of the specified strains. These rDNA-based data corroborate the CFU-based findings in Fig 3.
